# Predicting regulators of epithelial cell state through regularized regression analysis of single cell multiomic sequencing

**DOI:** 10.1101/2022.12.29.522232

**Authors:** Nicolas Ledru, Parker C. Wilson, Yoshiharu Muto, Yasuhiro Yoshimura, Haojia Wu, Amish Asthana, Stefan G. Tullius, Sushrut S. Waikar, Giuseppe Orlando, Benjamin D. Humphreys

## Abstract

Chronic disease processes are marked by cell-specific transcriptomic and epigenomic changes. Single nucleus joint RNA- and ATAC-seq offers an opportunity to study the gene regulatory networks underpinning these changes in order to identify key regulatory drivers. We developed a regularized regression approach, RENIN, (**Re**gulatory **N**etwork **In**ference) to construct genome-wide parametric gene regulatory networks using multiomic datasets. We generated a single nucleus multiomic dataset from seven adult human kidney biopsies and applied RENIN to study drivers of a failed injury response associated with kidney disease. We demonstrate that RENIN is highly effective tool at predicting key *cis-* and *trans-*regulatory elements.

## Introduction

Transcriptional control is coordinated through the binding of *trans*-acting transcription factors (TFs) to DNA motifs present in short, nucleosome-free genomic *cis*-regulatory elements (CREs)(1). These CREs allow the binding of different sets of transcription factors and subsequent recruitment of co-activators and transcription complex factors through dynamic regulatory activity and looping interactions with target gene promoters(2–4). Misregulation of gene transcription has the potential to drive disease (1,5–7), so identifying disease-associated CREs and their gene targets, as well as the key set of transcription factors coordinating this regulatory action, will improve our understanding of transcriptional regulation in disease and reveal novel therapeutic strategies.

Single cell sequencing has been successfully applied to characterize the transcriptional changes associated with specific cell types and cell states in a variety of human kidney diseases (8–15). Several current challenges limit our ability to leverage these datasets for the identification of regulatory factors coordinating gene expression changes. For example, it is difficult to prioritize the handful of transcription factors responsible for a particular cell process among the hundreds of cell-specific TFs and cognate motifs that can be detected(16). New tools such as single-cell regulatory network inference and clustering (SCENIC) identify coexpression modules from snRNA-seq datasets using TF binding motifs located near the transcription start sites of putative target genes in order to identify the TFs driving module regulation (17). The majority of regulatory elements are located in intergenic or intronic regions distal to the promoter, however (6,18,19), and these distal regulatory elements cannot be detected in snRNA-seq datasets. Methods to identify relevant CREs and TFs for any gene of interest across the genome are needed.

Single cell multiomic sequencing generates joint RNA- and ATAC-sequencing data from the same cell. In particular, it provides the ability to couple regulatory element activity to gene transcription(20). We hypothesized that this could be used to identify the regulators of injury and repair responses driving disease states, where a cell-specific healthy-to-disease trajectory with associated transcriptional changes is known. We developed a computational tool, RENIN (**Re**gulatory **N**etwork **In**ference) to construct genome-wide parametric gene regulatory networks that generates predicted weights for both CREs and TFs to rank and prioritize the most important regulatory elements. We built on current approaches linking individual CREs to individual genes with joint single cell RNA- and ATAC-seq(20) and a gradient boosting-based approach(21) by introducing the adaptive elastic-net estimator for gene regulatory network model training to increase model sparsity and thus improve regulatory element selection(22). We hypothesized that our parametric design would improve regulator prioritization when attempting to identify critical regulators of a given disease process. We then used motif data to identify transcription factors binding to these putative CREs and introduced a second step to correlate expression of predicted binding transcription factors (TFs) to expression of target genes. RENIN therefore simultaneously predicts CRE- and TF-gene regulatory interactions from single cell multiomic datasets to predict detailed genome-wide gene regulatory networks.

To validate our approach, we generated single nucleus multiomic datasets from healthy human adult kidney samples and applied RENIN to identify regulators of a proximal tubule (PT) state that may drive fibrosis and inflammation, thereby increasing risk for chronic kidney disease (CKD). In acute kidney injury (AKI), PT cells adopt a dedifferentiated and proliferative phenotype, providing the kidney with some regenerative capacity post-injury(23). These cells are marked by phosphatidylserine receptor KIM1 (*HAVCR1*) expression, which binds apoptotic cell fragments to clear debris from the tubular lumen(24). During the repair process, a small proportion of injured PT cells fail to undergo full repair and restoration of a healthy PT phenotype(9,25,26). These so-called failed repair PT (FR-PT) cells, first described in a mouse model of AKI, are likely to be important clinically, because they take on a proinflammatory, profibrotic, senescent-associated secretory pathway (SASP) phenotype, that may increase risk for CKD development (9,10,26).

Mouse FR-PT cells closely resemble a human proximal tubule state, PT_VCAM1, that is marked by VCAM1 expression and has been identified in healthy, non-AKI, human kidney(14,27). The PT_VCAM1 population increases with age and in diabetic kidney disease (10,14,28). Although FR-PT arise after acute injury, the transcriptionally similar PT_VCAM1 state is present even in healthy kidneys and we hypothesize it represents a “wear and tear” or injury *in situ* state, even in the absence of clinical AKI. We therefore asked whether RENIN could predict the regulatory elements driving formation of PT_VCAM1 cells in our dataset. We used RENIN to identify cell type-defining regulatory elements for all major kidney cell types, as well as those driving the differential expression changes seen between healthy PT and PT_VCAM1 populations. Our parametric approach significantly enriched for disease-relevant regulatory elements and had improved recall of cultured proximal tubule cell (RPTEC) open enhancer regions. Partitioned heritability showed that RENIN had increased enrichment of kidney-relevant traits compared to Signac’s LinkPeaks and DIRECT-NET (20,21). We used the model-trained parameters to prioritize key TFs driving the healthy to FR transition and selected a high-yield TF for orthogonal validation. We confirmed that one predicted driver of the failed repair state, NFAT5, promotes an *in vitro* phenotype resembling FR-PTCs, demonstrating the utility of our approach in the discovery of pathogenic signaling pathways. These findings demonstrate that RENIN may be applied to predict and characterize transcriptional regulatory mechanisms in disease.

## Results

### Simultaneous single-nucleus transcriptional and chromatin accessibility profiling of the adult human kidney resolves high-quality cell type-specific profiles

We performed single nucleus multiomic sequencing on seven control adult kidney samples. Five samples were nephrectomy-derived, and two samples were back-bench biopsies taken from pre-transplant healthy kidneys. Patient samples were heterogenous in age, sex, and race: median 64 years old (range: 45-76) 2 male and 5 female, and 3 Black and 4 White. Five of these patients had normal kidney function and two of them had advanced chronic kidney disease (Supplemental Table 1). Histologic scoring of interstitial fibrosis and tubular atrophy (IFTA) was 1-10% for all but one sample which had 10-20% IFTA in the setting of a serum creatinine of 4.73 mg/dL. Nuclei isolation was performed for each sample one at a time, followed by library generation and sequencing. Expression profiles were corrected to remove ambient RNA contamination with CellBender, predicted doublets were removed, then batch correction was performed with Harmony on both the split RNA-seq and ATAC-seq portions of the dataset(29–32). A total of 50,768 nuclei were annotated with cell types following quality control filtering. Among these nuclei, 29,758 genes and 193,787 accessible chromatin regions were identified. 150,237, or 77.5%, of peaks overlapped with a previously published snATAC-seq dataset(28), confirming quality (Supplemental Figure 1A). Overall, correlation between snRNA-seq expression and snATAC-seq gene activity was 0.552 (p <2e-16), but was higher between shared variable features in the RNA and ATAC datasets (r = 0.749, p < 2e-16; Supplemental Figure 1B). To compare the value added by single cell multiome vs. performing snRNA-seq and snATAC-seq on separate samples, we tested Seurat’s cell transfer annotation method(33) on our multiome data as though it was generated separately. Performance was high for certain cell types, such as the loop of Henle (LH), but low for other cell types, including podocytes (POD) and mesangial cells (MES). For example, 93.8% of the mesangial cells were predicted to be endothelial cells (Supplemental Figure 1C). These findings suggest multiome sequencing improves the ability to resolve cell types and states without sacrificing gene and peak detection sensitivity.

We performed weighted nearest neighbor clustering, an approach that learns relative cell type specific mRNAs and chromatin peaks to define cell types and states in the dataset (34). All major renal cortex cell types described in previous studies were successfully identified by annotating clusters based on snRNA-seq marker expression: proximal convoluted tubule (PCT), proximal straight tubule (PST), *KIM1+* proximal tubule (KIM1+ PT), parietal epithelial cells (PEC), loop of Henle (LH), distal tubule and collecting tubule (DCT, DCT/CNT, CNT), collecting duct (PC, ICA, ICB), podocytes (POD), endothelium (ENDO), mesangium (MES), fibroblasts (FIB), and immune cells (Immune) (Figure 1A). Notably, our multiomic clustering allowed separate annotation of PCT and PST clusters, which had been previously difficult to resolve using snRNA-seq alone(28). Cell type specific marker expression and gene activity, measured by aggregating accessible peaks within the gene body and promoter, was broadly similar between datasets (Figure 1B).

**Figure 1.**
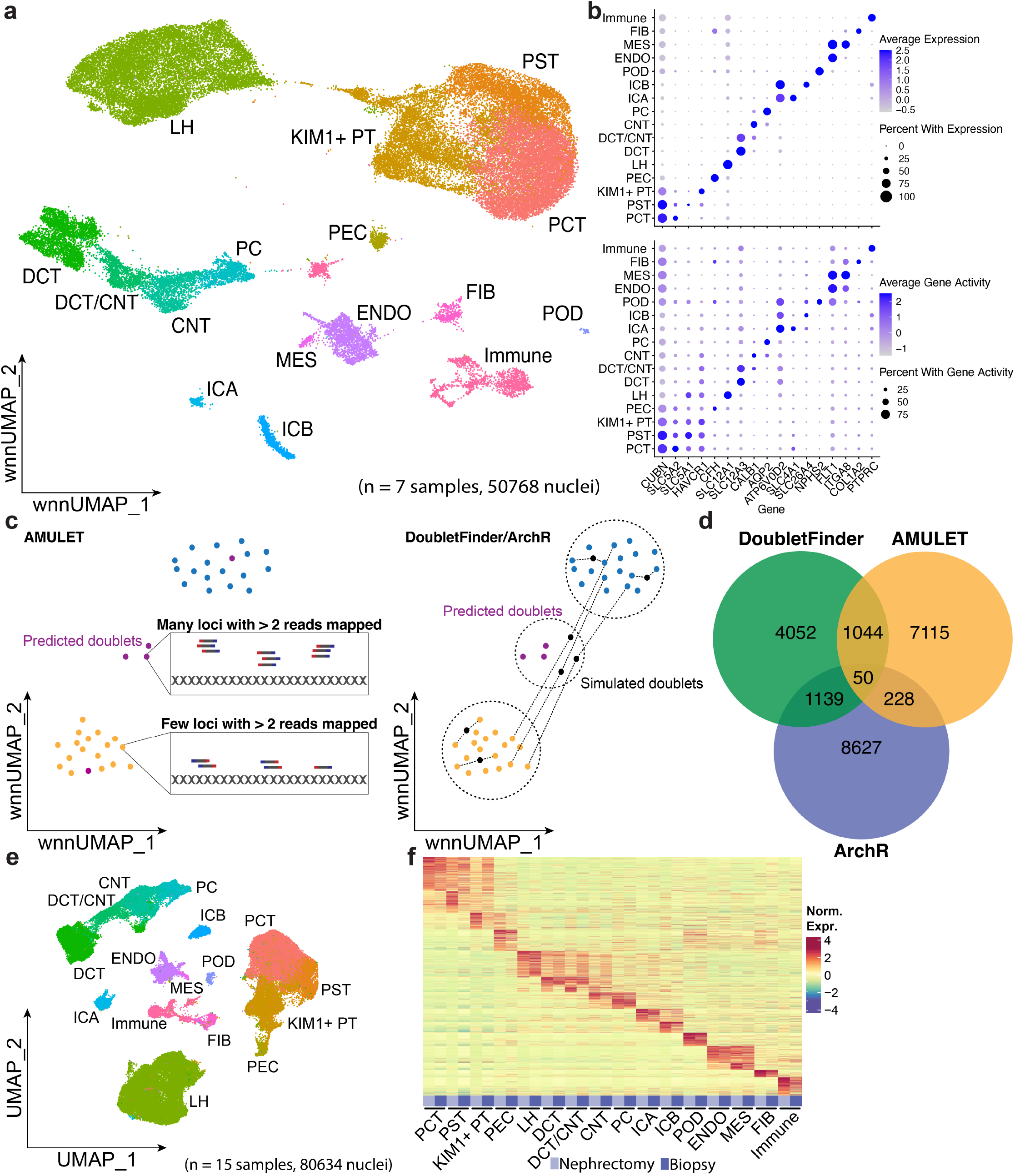
Simultaneous single nucleus multiomic RNA-seq and snATAC-seq of adult human kidney. **a**. WNN UMAP plot of multiome dataset prepared from 7 samples and totaling 50,768 nuclei. PST, proximal straight tubule; PCT, proximal convoluted tubule; KIM1+ PT, KIM1-expressing injured/failed repair proximal tubule; PEC, parietal epithelial cell; LH, loop of Henle; DCT, distal convoluted tubule; CNT, connecting tubule; PC, principal cell; ICA, intercalated alpha; ICB, intercalated beta; POD, podocyte; ENDO, endothelial; MES, mesangial; FIB, fibroblast. **b**. Above, RNA expression of cell type markers by cell type; below, gene activity, of cell type markers by cell type. Gene activity calculated by aggregating promoter and gene body peaks in snATAC-seq dataset. **c**. Left, cartoon of doublet finding approach used by AMULET. Nuclei have two copies of a given genomic locus, so barcodes with many loci with greater than 2 reads mapped are predicted to be doublets. Right, cartoon of doublet finding approach used by DoubletFinder and ArchR. Doublets are generated by averaging the expression or chromatin accessibility profiles of two barcodes in the dataset. Barcodes are predicted to be doublets by the proportion of neighboring barcodes that are simulated doublets versus original barcodes. **d**. Venn diagram of predicted doublets by each algorithm. **e**. UMAP plot of aggregate snRNA-seq dataset generated from a total of 15 samples (5 from living donor biopsies from 3 individual donors and 10 from nephrectomy tissue), containing 91,337 nuclei. **f**. Heatmap of cell type marker expression for each cell type by sample type—nephrectomy or biopsy.

### Doublet calling algorithms identify non-overlapping putative doublets

Droplet-based microfluidic methods commonly result in multiplets when more than one nucleus is encapsulated within the same droplet(30). When this happens, nucleotide fragments from multiple nuclei are tagged with the same barcode and thus appear as the same individual nucleus in downstream clustering, cell type annotation, and analysis. Several bioinformatic approaches have been developed to identify which barcodes are multiplets, including DoubletFinder, AMULET, and ArchR(30,31,35). DoubletFinder, using scRNA-seq data, and ArchR, using scATAC-seq data, both simulate artificial doublets by averaging profiles from pairs of randomly selected barcodes. Barcodes are identified as doublets, rather than single nuclei, by a high proportion of artificial doublets to dataset barcodes in neighboring points in a low dimensional projection of the dataset. Unlike DoubletFinder and ArchR, AMULET’s algorithm takes advantage of the fact that each individual nucleus contains only two copies of each genomic locus. In the absence of copy number variation, the maximum number of reads mapping to a given region should be two in a scATAC-seq dataset. AMULET determines the number of regions with greater than two overlapping fragments and identifies as multiplets barcodes with higher-than-expected numbers of these regions (Figure 1C). It is difficult and costly to test these approaches with biological “ground-truth” datasets, so simulated datasets have been used to compare performance of doublet identification methods(36). We reasoned that simultaneous multimodal single nucleus sequencing provides a unique opportunity to cross-assess doublet calling algorithms using data from different modalities. We performed doublet calling with AMULET and generated artificial doublets from the same pairs of starting nuclei, using either the RNA modality for DoubletFinder or the ATAC modality for ArchR. Unexpectedly, while overlap between the three tested methods was greater than random chance, it was still very low (<20%), consistent with previously reported ArchR-AMULET differences (Figure 1D) (31). AMULET-derived doublets averaged high ATAC unique molecular identifier (UMI) counts, concordant with the algorithm’s approach, whereas ArchR-doublets averaged low ATAC UMI counts (Supplemental Figure 2A). Unlike ArchR or DoubletFinder, AMULET can call homotypic doublets (Supplemental Figure 2B), so we removed AMULET-predicted doublets. Furthermore, when comparing doublets detected by DoubletFinder vs. ArchR, doublets predicted by ArchR were evenly spread throughout all clusters whereas those predicted by DoubletFinder were enriched between and on the edges of clusters, where we would expect them to be located (Supplemental Figure 2B). For this reason, we additionally removed DoubletFinder-predicted doublets.

### Partial nephrectomy kidney samples are similar to live donor samples and both contain FR-PT cells

FR-PTC cells have been identified in healthy human kidney samples derived from tumor-adjacent tissue after partial nephrectomy(14,28). We hypothesize that FR-PTC in apparently healthy kidneys is attributable to subacute injury or stress sustained over a patient’s lifetime, leading to accumulation of FR-PTC cells and a fibrosis-promoting phenotype. An alternative hypothesis is that tumor mass effect in partial nephrectomy samples elicits an acute injury stimulus that drives FR-PTC accumulation. Since living donor kidney biopsies are the gold standard source of ‘healthy’ human kidney tissue, we would predict that these samples would not contain FR-PTC if the alternative hypothesis were true.

We generated snRNA-seq and snATAC-seq (separately, not multiome) libraries from three additional human donors to increase sensitivity. We also added 5 previously published snRNA-seq and snATAC-seq datasets from nephrectomy samples for the same reason. We then compared nephrectomy-derived samples (n = 10) to living donor kidney biopsy samples (n = 5 from 3 individual donors) to test this alternative hypothesis. Aggregation of these samples yielded 80,634 cells from snRNA-seq and 120,679 cells from snATAC-seq. We could still clearly detect *KIM1*+/*VCAM1*+ FR proximal tubule cells even when clustering cells solely from living donor kidney biopsies, consistent with recent analysis of living donor kidneys (37), indicating that tumor mass effect alone does not explain FR-PTC accumulation (Figure 1E, Supplemental Figure 3A-D). More broadly, cell type-specific RNA expression and ATAC accessibility profiles were similar between living donor kidney biopsies and nephrectomy samples (Figure 1F, Supplemental Figure 3E). KIM1+ PT cells derived from either tissue source expressed a set of genes that were not expressed in healthy PT cells. These findings suggest there may be minor differences between partial nephrectomy and living donor kidney-derived cell populations, like *KIM1*+/*VCAM1+* PT, which may be affected by the lower number of living donor samples included. Detection of FR-PT cells in living donor-derived samples supports our usage of both tissue sources in analyzing the FR-PTC multiome.

### Regulatory network inference with adaptive elastic-net model

A major advantage of simultaneous RNA-seq and ATAC-seq measurements from the same cell by identifying peak-gene pairs with correlated accessibility and expression. This correlational analysis generates hypotheses for CREs that are forming distal looping interactions with target gene promoters. Currently available tools such as Signac’s LinkPeaks function employ a univariable regression model to correlate chromatin accessibility to gene expression. However, testing peak-gene pairs individually has some limitations. In snATAC-seq datasets, chromatin accessibility values between nearby peaks are not always fully independent, a feature that has been leveraged previously to assemble networks of *cis*-coaccessible peaks(38). For most genes, multiple CREs coordinate to regulate expression (Supplemental Figure 4), so a multivariable regression model is more appropriate for modeling the influence of CRE accessibility on gene expression. Additionally, it stands to reason that adjusting for the influence of all CREs within a regulatory network facilitates more accurate selection of key regulatory elements with lower false discovery rates relative to a univariable modeling approach for a given gene’s expression.

However, single cell multiome datasets have some characteristics that may impede model performance. The sparse and high-dimensional nature of single cell datasets makes it difficult to select a minimal set of regulatory elements for a given gene, with hundreds of possible peaks and TFs that may target any given gene. Within gene regulatory networks, groups of genes can be regulated by the same regulatory elements, and the resulting collinearities increase the difficulty of selecting the most important regulatory elements. The adaptive elastic-net has several features that are well-suited for single cell datasets(22). First, model complexity is penalized, which enables variable selection in high-dimensional datasets. An adaptive L1 penalization reduces false positives in selecting regulatory elements for a given gene. Importantly, the adaptive L1 penalty addresses two critical weaknesses with a common choice for modeling this kind of dataset, the lasso estimator—instability in variable selection and lack of oracle property—by introducing adaptive weights for penalizing each coefficient (39). An additional L2 penalization term increases model stability in high dimensional datasets with collinear predictor variables, which is a defining feature of single cell datasets. Finally, parametric model construction allows for ranking of CREs, thus improving yield when selecting potential key CREs for biological follow-up and validation. In relation to other options, the adaptive elastic-net configuration of our model is optimal for identification of *cis*-regulatory elements regulating a target gene.

We reasoned that TF motifs in candidate CRE provide additional information beyond chromatin accessibility alone. Therefore our model introduces a second adaptive elastic-net step, in which putative *trans*-regulatory elements (TFs) are linked to a gene through TF binding motif information, then expression of these TFs is used to predict the target gene’s expression (Figure 2). TFs binding candidate CREs, but with low and/or uncorrelated expression, are pruned, leaving fewer but higher quality candidate TFs for biological validation. With this two-step approach, our model can select both key CREs and key TFs to construct gene regulatory networks for a given gene of interest.

**Figure 2.**
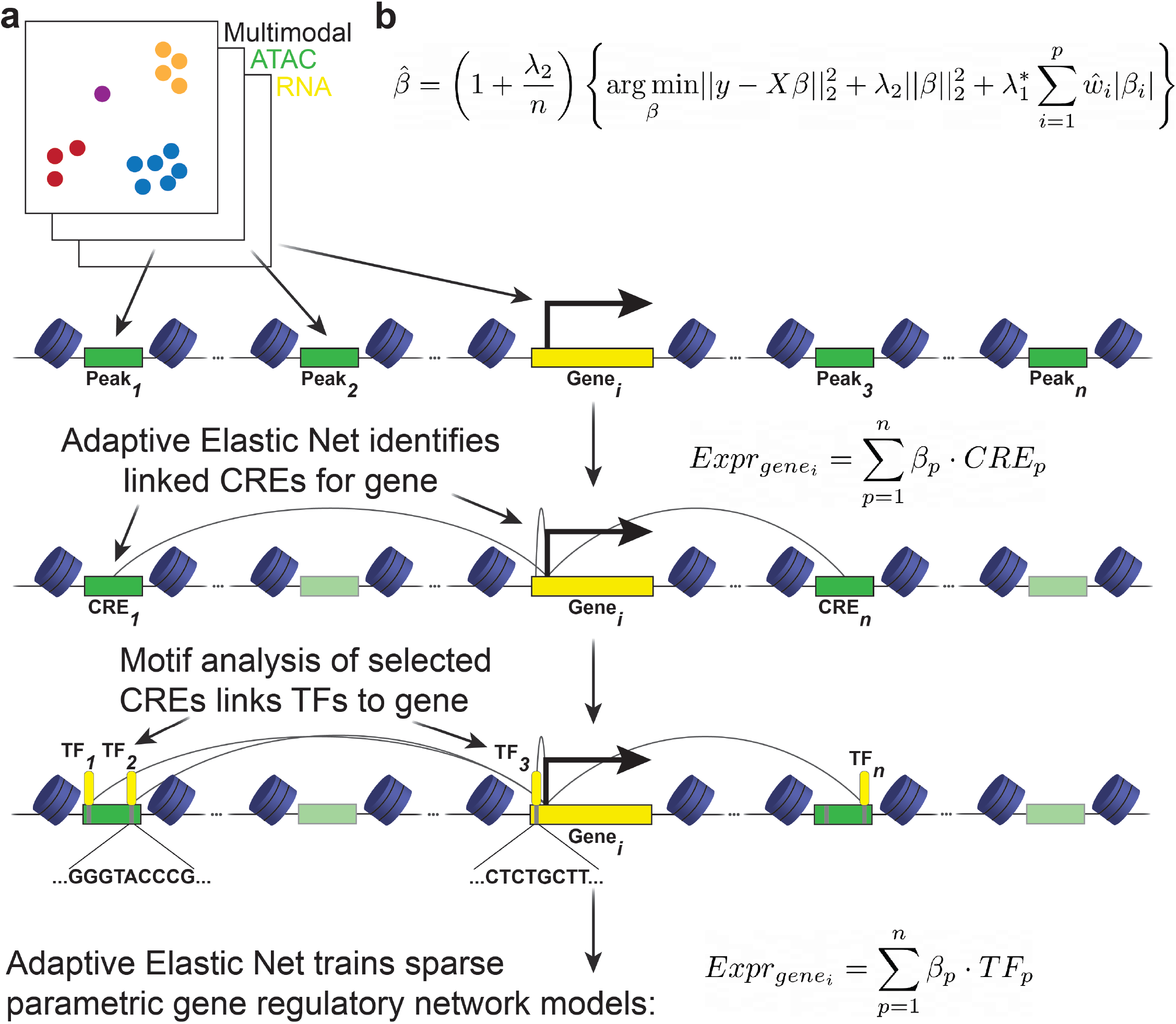
Overview of model design. **a**. Clustered multiome dataset contains chromatin accessibility and gene expression profiles for each nucleus. The model’s first step is to learn gene expression predicted by accessibility of peaks within 500kbp of the gene TSS. This step identifies CREs as peaks with accessibility changes correlated with target gene expression. The second step annotates peaks with potential binding TFs by scanning for TF motifs. TFs with predicted motifs in predicted CREs are aggregated as putative regulatory TFs for a target gene. The third step is a repeated training step in which the model learns gene expression predicted by expression of TFs selected in the second step. This step identifies putative regulatory TFs based on the correlation between target gene and TF expression in the multiome dataset. For both learning steps, an adaptive elastic-net regression model is used. **b**. The adaptive elastic-net estimator used for model fitting.

### Cell type-specific *cis*-regulatory elements identified with RENIN

We tested the ability of RENIN to select functional CREs by using it to identify CREs regulating cell type-specific gene expression. We identified an average of 893 (range: 612-1304) cell-specific genes per cell type (Supplemental Data 1). For each of these genes, our model considered all peaks within 500kbp of the TSS. The model identified 32,304 total unique CREs, with an average of 4263 CREs predicted to be active (range: 3016-11006) per cell type, highlighting the utility of studying cell types that are sampled at different rates in single cell datasets (Figure 3A, Supplemental Data 2). Of the 6,105 modeled genes, 5,038 had at least one linked CRE, suggesting that CRE accessibility is an important regulatory mechanism in human adult kidney. Most (27,934) CREs had one target gene, with 4,370 having two or more targets (Supplemental Figure 5A). We calculated a CRE’s regulatory score on a given target as the model-trained coefficient of that CRE, weighted by its average accessibility. Out of the total 62,255 predicted CRE-gene links, 13,489 links had negative predicted regulatory scores, suggesting this approach may also be useful in the study of silencers. Predicted regulatory weights of CREs annotated as promoters (peaks within 2kbp of the target gene TSS) were higher on average than either gene body or intergenic peaks, suggesting our approach assigns quantitative regulatory scores appropriately (Figure 3B). However, beyond the promoter, regulatory scores for intergenic and gene body CREs did not decline with distance, suggesting that RENIN identifies functional CREs rather than regions that are accessible simply due to proximity to an open and transcribed TSS.

**Figure 3.**
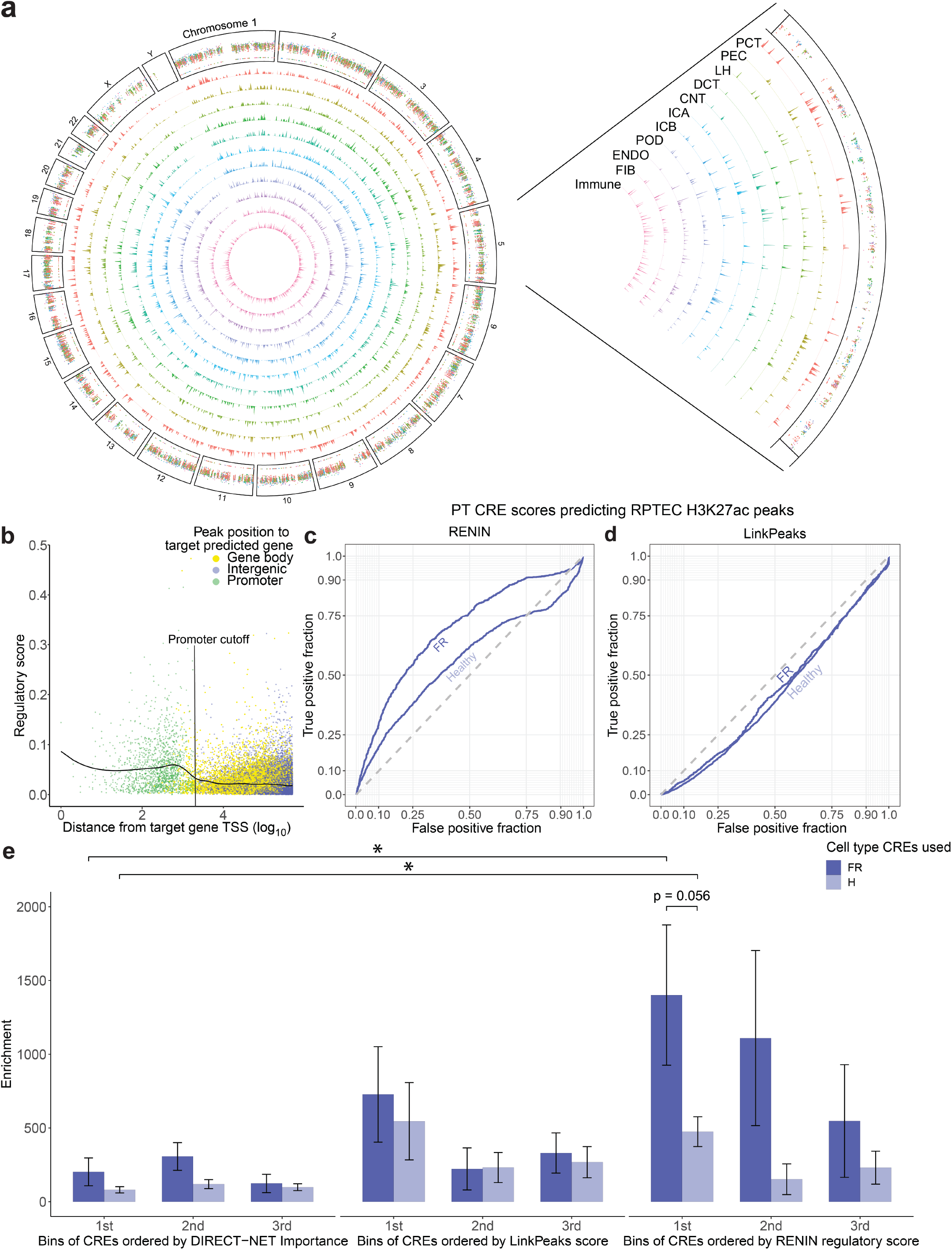
Cell type CREs identified with RENIN. **a**. Plot of CREs regulating cell type marker genes across genome by cell type. Chromosome 5 is expanded for visualization. **b**. Predicted CRE regulatory score, summed absolute value of CRE coefficients, by distance from target gene transcription start site (TSS). Peaks falling within target gene promoter are colored green, peaks within target gene body are yellow, and intergenic peaks are purple. Black line is smooth spline curve calculated on data points to illustrate trend over distance. **c**. ROC curve calculated for RENIN-predicted healthy-FR PT CREs against RPTEC H3K27ac peaks identified with CUT&RUN. FR (AUC = 0.694, CREs predicted to regulate a marker gene of the KIM1+ cluster; Healthy (AUC = 0.570), CREs predicted to regulate a marker gene of the PCT and/or PST cluster. Dotted line indicates random performance (AUC = 0.5). **d**. ROC curve calculated for LinkPeaks-predicted healthy-FR PT CREs against same RPTEC H3K27ac peak set as in (**c**). AUCs were 0.435 (FR) and 0.422 (Healthy). **e**. Comparison of RENIN, LinkPeaks, and DIRECT-NET by enrichment of partitioned heritability of CKD in model-predicted healthy (PCT + PST) and FR (failed repair—KIM1+ PT) CREs. Statistical significance determined by two-tailed t-test of difference between enrichment, p < 0.05.

In order to validate RENIN’s CRE predictions, we performed Cleavage Under Targets and Release Using Nuclease (CUT&RUN) sequencing on cultured RPTECs(40). CUT&RUN uses histone modification-specific antibodies to target MNase cleavage to nearby DNA, allowing localization of histone modification peaks in the genome. We performed CUT&RUN for two histone modifications marking active chromatin: H3K27ac and H3K4me3. Peaks were called with MACS2 to generate ground-truth PT CRE sets to assess RENIN’s performance. We constructed receiver operating characteristic (ROC) curves of RENIN’s ability to recover CUT&RUN peaks. FR-PT and healthy (PCT and PST) PT predictions both performed better than random chance for recovery of H3K4me3 peaks: area under the curve (AUC) for PT_VCAM1 predictions was 0.696 and 0.607 for healthy PT (Supplemental Figure 5B). RPTECs cultured on plastic partially recapitulate the FR phenotype(28). The higher AUC of PT_VCAM1 predictions suggests that cultured RPTECs may be closer to a FR than healthy phenotype. This difference is increased when using the H3K27ac peak set: PT_VCAM1 AUC is 0.694 and healthy AUC is 0.570 (Figure 3C). Since H3K27ac may be more of a marker of enhancer accessibility than H3K4me3(41), RENIN may be particularly useful for identifying distal CREs, which may vary more in accessibility depending on cell states whereas gene body accessibility is more static. Furthermore, the sparsity of snATAC-seq relative to bulk ATAC-seq may limit peak calling, so increased sequencing depth may improve recall as well.

We next compared RENIN with current state-of-the-art approaches including Signac’s LinkPeaks and DIRECT-NET(21). The univariable approach taken by LinkPeaks resulted in AUCs of 0.435 and 0.422 for PT_VCAM1 and healthy CREs, respectively (Supplemental Figure 5C). The sub-0.500 AUC in comparison with the AUC computed with RENIN-predicted CREs confirms that a multivariable approach is best suited for the study of chromatin accessibility due to correlations in accessibility between peaks. RENIN also outperformed DIRECT-NET, which predicted PT_VCAM1 and healthy CREs resulting in 0.631 and 0.621 AUCs, respectively, using the H3K27ac peak set (Supplemental Figure 5D-E). Notably, the PT_VCAM1-healthy AUC difference decreased when using DIRECT-NET predictions. Since RPTECs appear to have an expression profile resembling the FR-PT/PT_VCAM1 state, RENIN may have improved resolution between cell states relevant in disease.

We confirmed RENIN’s ability to identify important CREs by partitioning heritability of kidney related GWAS loci into predicted cell type CREs. Proximal tubule cells have been previously reported to enrich for heritability of CKD and eGFR(14). Cell type partitioning is closely related to cell-specific function and gene expression. We hypothesized that RENIN-identified CREs of cell type-specific genes would enrich for heritability of these traits(42). We found statistically significant enrichment of both traits in both healthy and PT_VCAM1 CREs, supporting RENIN’s utility in identifying functional CREs for traits with a genetic component (Figure 3E, Supplemental Figure 6). We binned healthy and PT_VCAM1 CREs by decreasing regulatory score (1^st^ bin to 3^rd^ bin, splitting the CRE set into equally sized thirds) and calculated partitioned heritability into each bin, finding a trend of increasing enrichment of CKD heritability(42) with increasing PT_VCAM1 CRE score. We also binned DIRECT-NET-identified CREs by Importance score and LinkPeaks-identified CREs by score. RENIN predictions had a statistically significant increase in enrichment relative to DIRECT-NET and a trending increase relative to LinkPeaks. Interestingly, the top PT_VCAM1 CRE bin had statistically significant higher enrichment of both tested traits than the top healthy CRE bin (Figure 3E, Supplemental Figure 6). We also observed a trending decrease in enrichment from the 1^st^ to 3^rd^ bin for both PT_VCAM1 CREs (p = 0.162) and healthy CREs (p = .106), suggesting the parametric ranking may prioritize CKD-relevant CREs. Taken together, these findings validate our approach to enrich for important regulatory regions and highlight the potential relevance of the PT_VCAM1 state in renal disease and declining kidney function.

### Cell type-driving TFs are highlighted with RENIN

We reasoned that a CRE predicted to regulate a nearby cell type-specific gene in *cis* should contain motifs for TFs that regulate that gene. We used the filtered cisBP motif database provided with the chromVAR package to annotate each CRE and gene promoter region with predicted TF binding sites(16). We attempted to prune this initial list to enrich for cell type-defining TFs. For each cell type-specific gene, the list of TFs with at least one predicted binding site within the CRE and promoter peak set comprised the initial list of putative TFs used for RENIN’s second step.

Next, we calculated cell type-specific regulatory scores for each TF by summing their regulatory coefficients weighted by their mean expression in each cell type. This allowed us to identify cell type-defining TFs for each renal cell type that replicated known biology (Figure 4A). For example, WT1, TCF21, and MAFB are known podocyte-specific TFs(43–45) and were identified as key podocyte TFs. PPARA and HNF4A, both well-characterized as driving PT differentiation and function(46,47), were top key proximal convoluted and straight tubule TFs. In contrast, PPARA and HNF4A had lower regulatory scores in FR-PTCs, consistent with their dedifferentiated PT phenotype. Notably, RENIN also performed well for rarer cell types: for example, candidate PEC-specific TFs include RFX2 and TP63. Intriguingly, evidence implicates PECs as progenitor cells and TP63 is a p53 family member implicated in regulation of stem cell state(48–51). We performed a head to head comparison of TF ranking between RENIN and ChromVAR(52), which is currently used to identify cell type-specific TFs from snATAC-seq sequence data alone. While chromVAR agreed with RENIN for some TFs, overall agreement was low (Figure 4B). As RENIN effectively uses both RNA expression and sufficient accessible binding sites to predict important cell type TFs, its predictions may be more accurate. For example, chromVAR predicted MAFB to be a significant TF in PCT, despite low RNA expression (Figure 4C). TFs ranked by RNA expression levels had better concordance with RENIN predictions than did chromVAR for selected known TFs, but was not uniformly equal for all selected TFs. RENIN’s regularized approach may therefore eliminate spurious cell type-driving candidate TFs by integrating motif and expression data.

**Figure 4.**
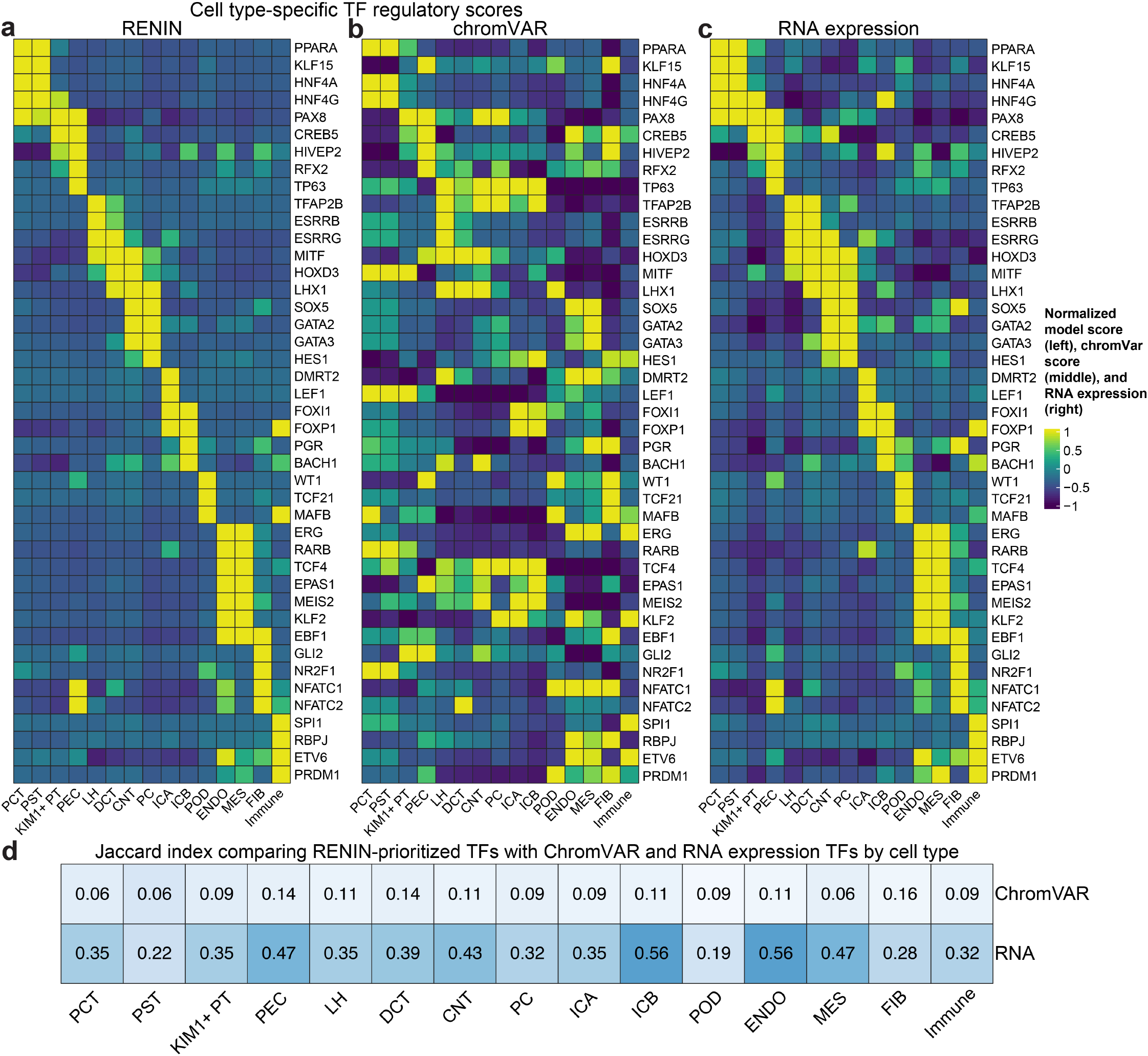
RENIN identifies cell type TFs. **a**. Heatmap of regulatory scores of select TFs by cell type. Regulatory scores calculated by summing regulatory coefficients of a TF on a cell type’s marker gene set and multiplying by mean expression of that TF in the given cell type. **b**. Heatmap of ChromVAR enrichment scores for select TFs by cell type. **c**. Heatmap of RNA expression for select TFs by cell type. **d**. Jaccard index between the top 25 TFs for each cell type predicted by RENIN versus ChromVAR enrichment or ranked expression of TFs.

To quantitate differences between these approaches, we calculated the Jaccard index (ranges from 0 to 1, denoting zero and perfect overlap, respectively) on the top 20 cell type TF predictions by RENIN and by ChromVAR. This revealed low overlap between the two approaches (Figure 4D). The Jaccard index was higher between RENIN predictions and the top 25 TFs per cell type based on their RNA expression alone, in line with our weighting of TFs by mean cell type expression, but RENIN still predicted new cell type TFs that would not have been prioritized with TF expression alone. For example, TCF12 and MAFB were not among the 25 highest expression TFs in podocytes, but were top RENIN predictions for podocytes. So, while RENIN’s regression approach still relies on high quality expression data, it may still be applicable to low expressed genes or rare cell types due to its design and integration of multiple modalities. These findings further demonstrate that RENIN is an effective tool to integrate both TF expression data and motif data to achieve more specific identification of key cell type-defining TFs, while motif enrichment approaches could be more applicable for identifying factors that affect CRE accessibility.

### CRE analysis of healthy-failed repair axis reveals coordinated regulatory element remodeling

Next, we applied RENIN to study the gene regulatory networks involved in the healthy to failed repair transition. We hypothesized that expression changes along this transition would allow RENIN to identify *cis*-and *trans-*regulatory elements driving the change in cell state. Using the proximal tubule subset of our dataset, we identified 1666 genes differentially expressed (adjusted p-value < 0.05) between healthy (PCT and PST) PT and FR-PTC (Supplemental Data 3). Of these, 1516 were non-mitochondrial and present in our EnsDb.Hsapiens.v86 genome annotation reference. RENIN’s first step identified 15,434 Healthy to FR-PTC CREs regulating 1,488 of 1,516 input genes, with an average of 11.4 CREs per differentially expressed gene (Supplemental Figure 7A). These findings strongly implicate distal chromatin remodeling in the development of the FR-PT state. On average, RENIN explained a proportion of variance of 0.417 for genes with at least one linked CRE, with a negative r^2^ for only 89 modeled genes (Supplemental Figure 7B). Thus RENIN was able to generate a list of candidate CREs implicated in the H-FR transition.

We then asked what factors might be driving changes in accessibility of these CREs, resulting in differential expression along the H-FR transition. We classified CREs as either healthy-(H CREs) or failed repair-promoting (FR CREs) based on their predicted enhancer or repressor role on differentially expressed genes; a CRE was labeled healthy-promoting if it enhanced a healthy-upregulated gene or repressed a FR-upregulated gene and vice-versa. We further hypothesized that motif enrichment might identify factors that were candidate pioneering TFs driving healthy- and failed-repair-promoting CRE accessibility. We calculated motif enrichment within H and FR CREs. The most enriched H CRE motifs were HNF4A/G, while the most enriched FR CRE motifs were AP-1 subunit motifs (Figure 5A). HNF4A/G and AP-1 subunit motifs were the least enriched in the other CRE peak set, suggesting they may be involved in adversarial processes promoting normal differentiation or fibrosis and a failed repair-associated phenotype. Stimuli altering the balance between the two may influence the likelihood of H-FR transition through the opening of new regulatory TF-FR gene links.

**Figure 5.**
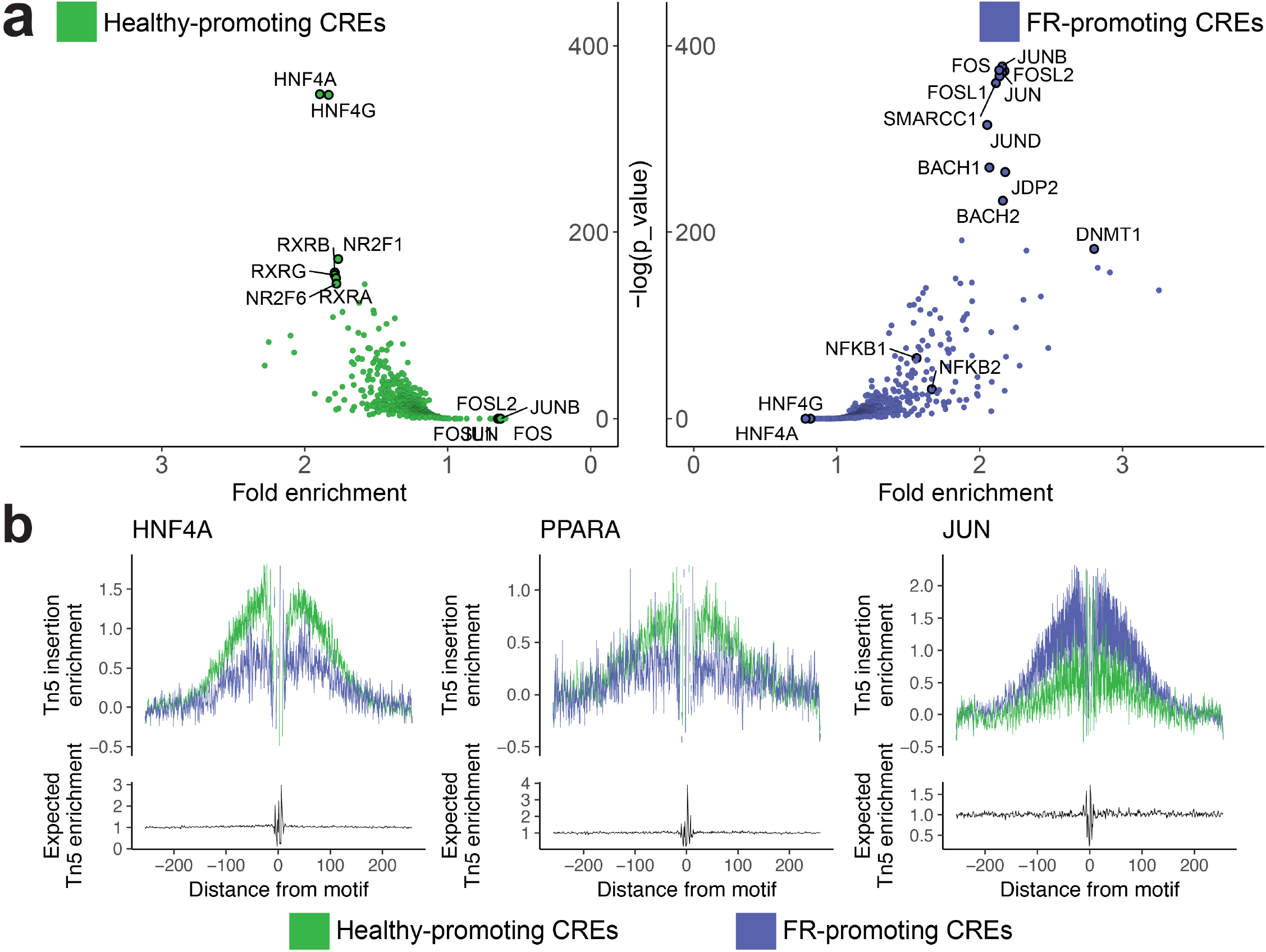
Analysis of CREs in healthy to failed repair PT transition. **a**. Motif analysis of CREs predicted to regulate differentially expressed genes between healthy PT (PCT and PST) and FR PT (KIM1+ PT) clusters. Left, healthy-promoting CREs were identified by predicted positive regulatory score on genes upregulated in healthy PT compared to FR PT or negative regulatory score on genes upregulated in FR PT compared to healthy PT. Right, FR-promoting CREs identified by positive score on genes upregulated in FR PT or negative score on genes upregulated in healthy PT. **b**. Footprinting analysis for HNF4A, left; JUN, middle; NFKB1, right. Tn5 insertion enrichment calculated around motifs present in healthy-promoting CREs (green) and motifs present in FR-promoting CREs (purple).

We also performed footprinting analysis to determine whether Tn5 insertion enrichment around motifs in H CREs and FR CREs was different. We found that Tn5 insertion was higher in regions adjacent to HNF4A and PPARA motifs in H CREs relative to those found in FR CREs, while Tn5 insertion was enriched near JUN motifs in FR CREs relative to those in H CREs (Figure 5B). NFKB1, previously identified as a likely factor in the H-FR transition(28), motifs also showed slight increased adjacent Tn5 enrichment relative to H CRE NFKB1 motifs, supporting the identification of these CREs as FR-associated or promoting (Supplemental Figure 7C). We also performed footprinting analysis for other TF motifs, and did not find the same differential insertion enrichment, demonstrating that this differential insertion enrichment is not due to non-specific differences in H and FR CRE accessibility (Supplemental Figure 7D). Our RENIN to chromVAR comparison suggested that enriched motifs within a target gene’s CRE sequences may not be the most important TFs regulating that gene. Instead, enrichment may provide information on overall peak accessibility or regulatiory activity. One possible mechanism affecting differences in binding in certain transcription factors may be methylation. We found that RENIN-predicted CREs were overlapped with CKD-associated methylated regions at a higher rate than the set of all called peaks in our dataset with an odds ratio of 1.322 (95% CI = 1.229 – 1.421, p = 2.138e-13 by Fisher’s exact test, Supplemental Data 4). Our dataset thus contains evidence that epigenetic remodeling of specific CREs, such as methylation, leading to altered TF binding is associated with the H-FR transition.

### Identification of healthy- and FR-promoting TFs with RENIN

The failed repair PT cell state is distinct from healthy proximal tubule cells and the aberrant phenotype is stable long after recovery from renal injury(9,10). We hypothesized that gene regulatory network activity in the proximal tubule could play a role in establishing and/or maintaining the failed repair population, and that there is a subset of key TFs regulating the transition. We constructed parametric gene regulatory networks for genes differentially expressed between healthy PTs and PT_VCAM1 cells in order to prioritize candidate TFs for this H-FR axis. We ranked TFs by the sum of their regulatory coefficients across all input genes, weighted by whether the target gene was FR-upregulated (negative score) or H-upregulated (positive score) and TF average expression (Figure 6A, Supplemental Data 5). Top healthy-promoting TFs included canonical PT TFs, HNF4A and PPARA, as well as other factors that have been demonstrated to be protective against kidney disease: ESRRG and RREB1(46,53–55). The top FR-promoting prediction was NFAT5, a TF that has been previously identified as upregulated following ischemia-reperfusion injury in a mouse model of AKI(10). Thus, RENIN prioritizes plausible TFs involved in the H-FR transition for further investigation. Moreover, while the initial list based solely on motif presence within a linked CRE averaged 392.2 TFs per gene (range: 18-602) model training resulted in sparse lists of putative TFs for each gene, (mean: 33.2, range: 4-125; difference in mean number of TFs by paired two-sided t-test p-value < 2.2e-16; Supplemental Figure 8A). This demonstrates an advantage of our approach: its ability to identify smaller subsets of higher importance TFs for further investigation.

**Figure 6.**
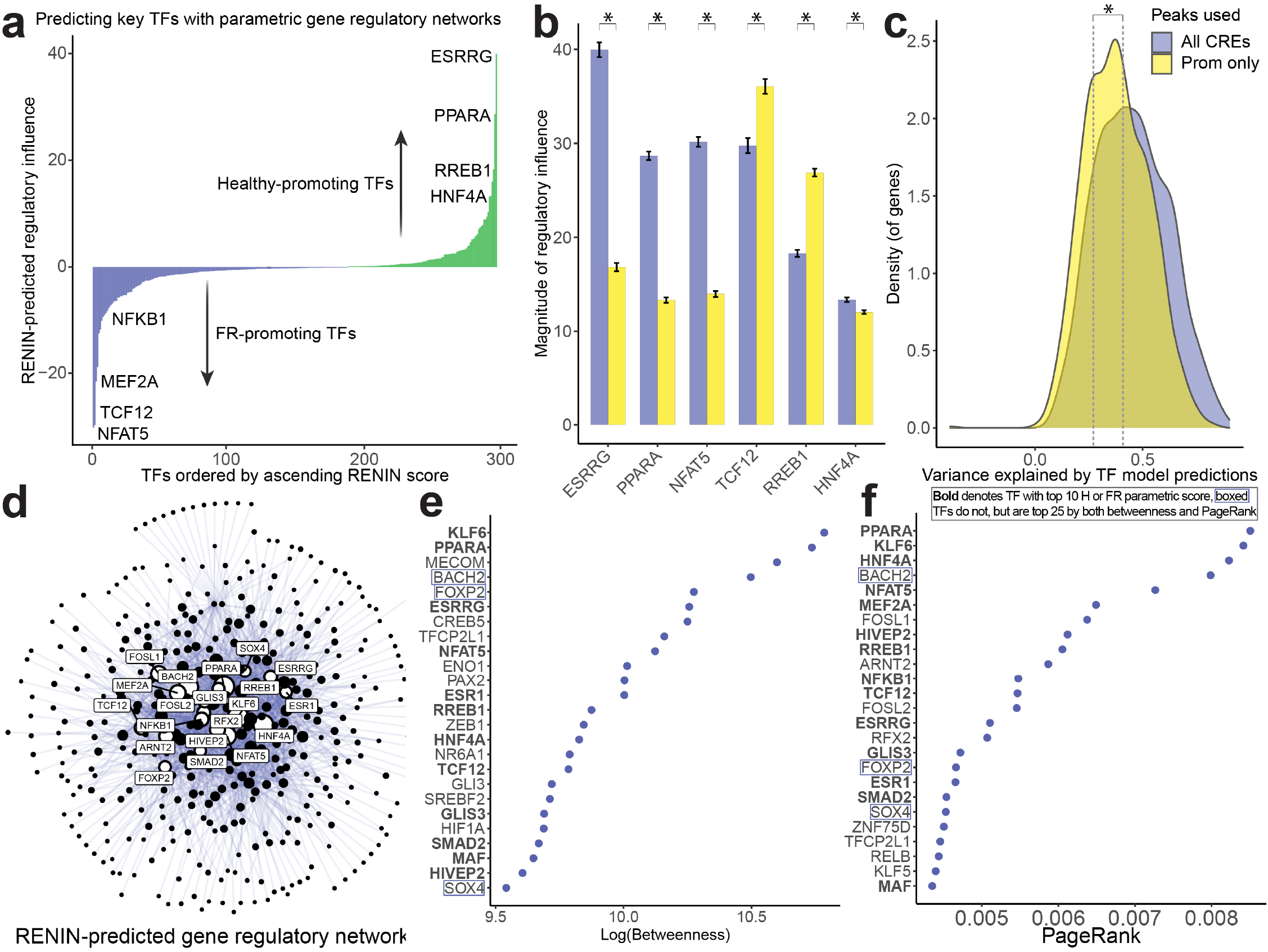
Predictions of key TFs involved in healthy to failed repair PT transition. **a**. TFs sorted by regulatory score, computed as the sum of predicted regulatory coefficients for healthy-FR PT DEGs multiplied by mean TF expression in PT (PCT, PST, KIM1+ PT) clusters. Negative scores indicate FR-promoting TFs— positive regulation of DEGs upregulated in FR PT or negative regulation of DEGs downregulated in FR-PT— and positive scores indicate H-promoting TFs—positive regulation of DEGs upregulated in H PT (PCT and PST) or negative regulation of DEGs downregulated in H PT. **b**. Regulatory scores calculated for selected TFs when model is trained with motifs from all linked CREs (purple) and from motifs only in target gene’s promoter region (yellow). Statistical significance determined by two-tailed t-test of the difference in mean scores, p < 0.05. **c**. Density plot of coefficients of determination of model predictions of DEG expression versus actual expression when training model with motifs from all linked CREs (purple, mean r^2^ = 0.45) and motifs only in target gene’s promoter region (yellow, mean r^2^ = 0.38). Statistical significance determined by two sample t-test, p < 2.2e-16. **d**. Graph visualization of gene regulatory networks predicted by RENIN. TF node size represents centrality of TFs computed by PageRank, top 25 TFs are labeled. **e**. Top 25 TFs ranked by betweenness. **f**. Top 20 TFs ranked by PageRank.

Given the coordinated chromatin remodeling previously described, we hypothesized that non-promoter CREs comprising a *cis*-coaccessibility network (CCAN) for a target gene have non-redundant regulatory influence due to the presence of additional TF binding sites that regulate the H-FR transition. We compared RENIN’s performance when using motifs present in CREs, as well as the promoter, to the promoter alone. We found that regulatory scores of ESRRG, PPARA, NFAT5, and HNF4A decreased, while TCF12 and RREB1 scores increased (Figure 6B). The addition of non-promoter CREs improves the proportion of variance explained by RENIN, measured by coefficient of determination (Figure 6C). In order to determine whether this was simply due to overfitting as a a result of the inclusion of more TFs, we compared the likelihoods of RENIN models when incorporating all CRE information versus promoter peak information alone by Akaike Information Criterion (AIC). For 1,323 genes that could be modeled with promoter-only and CCAN configurations, the vast majority—1,184 genes—were modeled more accurately with the distal CRE inclusion, and we calculated the mean ΔAIC to be −1293 (Supplemental Figure 8B, Supplemental Data 6). These findings strongly support the relevance of distal regulation, as the inclusion of the information they provide improves model fit. For the remaining 165 genes without any accessible promoter peaks using our reference annotation, RENIN was able to identify CREs and subsequently TFs with the inclusion of distal interactions. These results demonstrate the added utility of implementing the first step of RENIN to identify both distal and proximal CREs in improving accurate modeling of gene regulatory networks with single cell datasets.

Next, we assembled predicted gene regulatory networks into a directed graph representation (Figure 6D). We ranked TFs by two measures of centrality, betweenness and PageRank, to prioritize most central TFs in the H-FR gene regulatory landscape (Figure 6E-F, Supplemental Figure 8C). Top central TFs included similar predictions of TF importance—e.g. ESRRG, PPARA, HNF4A, RREB1, NFAT5, TCF12, and MEF2A were all within the top 20 central TFs by both measures. Simulated knockdown of 4 top predicted healthy-promoting TFs and simulated upregulation of 4 top predicted failed repair-promoting TFs caused simulated healthy PCT and PST cells to approach the KIM1+ PT cluster when plotted along the two top principal components, providing further computational evidence that these top predicted TFs are relevant in the H-FR transition (Supplemental Figure 9A). We designed RENIN to prioritize key TFs and the consistency between multiple methods of ranking supports its usage in identifying disease-relevant TFs for biological validation.

### *NFAT5* knockdown partially reverts failed repair phenotype

RENIN identified NFAT5 as the top TF driving the H-FR transition. We next used the RENIN-predicted NFAT5 gene regulatory network to simulate NFAT5 knockdown in FR-PT cells to predict the resulting effect on cell phenotype. Visualizing cells in the low dimensional PCA space computed by Seurat, simulated knockdown of NFAT5 in a sample of KIM1+ PT cells caused them to move towards healthy PCT and PST cells, consistent with NFAT5’s predicted FR-promoting effect (Supplemental Figure 9B). We then asked whether actual siRNA knockdown of NFAT5 in RPTECs would reduce the FR-PT phenotype in culture. We targeted *NFAT5* for siRNA knockdown and achieved a robust reduction in *NFAT5* mRNA levels, which was associated with decreased expression of the FR-PTC marker *VCAM1* (Figure 7A). Next, we performed bulk RNA-seq on *NFAT5* siRNA knockdown samples compared to control. We found reduced expression of fibrosis-associated genes including *TGFB1, TGFB1R, COL1A1*, and *COL4A1* (Figure 7B). We performed KEGG enrichment analysis on RENIN-predicted direct NFAT5 target genes and on the genes differentially expressed in *NFAT5* siRNA samples relative to non-targeting siRNA samples. RENIN predictions identified some shared enriched pathways with *NFAT5* knockdown-enriched pathways (Figure 7C). In *NFAT5* knockdown samples, we identified enrichment of ECM-receptor interaction, TGF-beta signaling, and other inflammation-related pathways (Figure 7D). Finally, we sought to confirm NFAT5 expression in adult human kidney cortex, as previous work on NFAT5 has focused on the medulla. Immunostaining for NFAT5 showed a patchy expression pattern, consistent with the hypothesis that FR-PTCs accumulate progressively in a scattered fashion over time. NFAT5 expression was anticorrelated with LTL staining, with LTL-low cells and tubules being highest in NFAT5 expression (Figure 7E). These results support the notion that NFAT5 promotes the failed repair PT cell state and suggest that NFAT5 could be a therapeutic target. They also demonstrate RENIN’s ability to identify key TFs for biological investigation.

**Figure 7.**
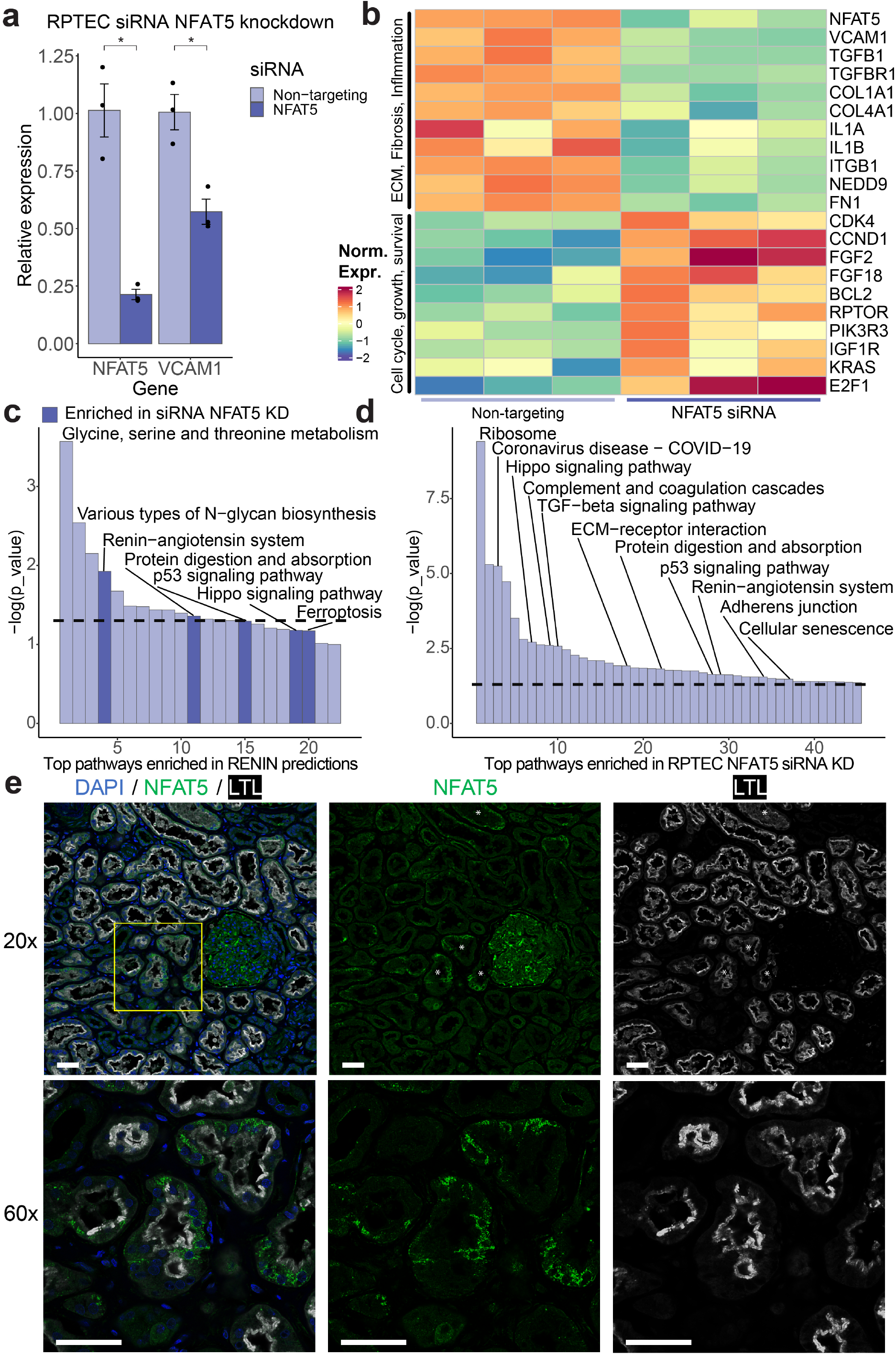
NFAT5 promotes FR expression phenotype in cultured RPTECs. **a**. Expression of *NFAT5* and *VCAM1* in cultured RPTECs treated with non-targeting (NT) or *NFAT5*-targeting siRNA. RNA levels measured by RT-qPCR and normalized to *GAPDH* expression. *NFAT5* siRNA-treated cells had 21% of the *NFAT5* RNA and 57% of the *VCAM1* RNA levels of non-targeting-siRNA-treated cells (p values of 0.017 and 0.012, respectively, unpaired 2-tailed t-test). **b**. Heatmap of select differentially expressed genes by RNA-seq in NT and *NFAT5* siRNA-treated RPTECs. **c**. KEGG pathway enrichment in RENIN-predicted NFAT5 target gene set. Dotted line corresponds to p value of 0.05. Dark purple bars correspond to pathways enriched in set of differentially expressed genes between NT and *NFAT5* siRNA-treated RPTECs. **d**. KEGG pathway enrichment in differentially expressed gene set between NT and *NFAT5* siRNA treatment. **e**. Immunofluorescent labeling of NFAT5 in adult human kidney. DAPI is a nucleus marker and LTL is an apical proximal tubule marker. Asterisks denote examples of tubules with low LTL intensity. Scale bars are 50 μm in length.

## Discussion

Profibrotic, proinflammatory PT cell states have been increasingly predicted to play a role in CKD development following incomplete PT repair following acute injury or during chronic progressive disease(10,14,28,56). Here we performed joint single nucleus RNA and ATAC sequencing on adult human kidney samples to study the PT_VCAM1 state in adult human kidney. We found that the presence of PT_VCAM1 was not exclusively explained by sample origin, as populations were also detected in biopsy-derived samples. These results support the use of nephrectomy samples as adequate control tissue in human kidney research, including for the study of the role of PT_VCAM1 cells in CKD.

We applied RENIN to our multiomic dataset to study both *cis*- and *trans*-regulatory machinery in the proximal tubule. We found that increasing predicted CRE regulatory influence was associated with increased enrichment of CKD heritability, further implicating PT_VCAM1 in CKD. Risk variants open in PT_VCAM1 CREs may exert their effects through amplifying profibrotic, proinflammatory signaling or increasing the accumulation and/or persistence of the PT_VCAM1 state. We also identified a small set of TFs predicted to regulate the healthy-PT_VCAM1 transition. Our model suggests targeting some candidate TFs such as NFAT5 and ESRRG may attenuate the PT_VCAM1 phenotype. For example, ESRRG may regulate mitochondrial function and metabolism in a protective capacity (46,53). NFAT5 has previously shown to play a protective role in AKI and against hypertonic stress(57). Our results suggest that chronic expression of NFAT5 may instead promote a proinflammatory phenotype in PT_VCAM1 cells.

Our multiomic dataset and catalog of cell type-specific regulatory elements will serve as a resource for kidney research but our modeling approach to comprehensively profile gene regulatory networks with multiomic single cell datasets should be more broadly applicable to other tissues as well. We found RENIN’s parametric design had great utility in highlighting both CREs and TFs that are relevant for our renal disease research interest through full integration of information from both expression and chromatin accessibility data. Our adaptive elastic-net-based approach minimizes risk of overfitting and misallocation of priority when selecting key regulatory elements relative to other methods. Greedy algorithms like gradient boosting approaches can suffer from overfitting (58) without careful titration of the learning rate, early stopping conditions, and other parameters, thus hindering regulator selection. In contrast, RENIN strives for model sparsity, which may reduce false positive rates for predicted regulators and explain the higher enrichment for risk variants and CUT&RUN peaks. Particularly in comparison to non-parametric approaches and decision tree approaches, the ability to effectively prioritize candidate regulatory factors for further investigation makes RENIN a highly effective computational tool for hypothesis generation.

RENIN identified cell type-specifying TFs in a manner more consistent with known biology than chromVAR did with a motif enrichment approach. It may be more accurate to leverage predicted regulatory relationships based on expression rather than motif enrichment, because the number of motifs may not gate TF regulatory activity. A TF may exert strong influence through few motifs, which allows RENIN to identify regulatory networks through expression information. Instead, motif enrichment in CREs may provide hypotheses for mechanisms regulating peak accessibility or overall regulatory activity. Furthermore, while RENIN may be limited by low mRNA expression, we found that it could still use the latent correlational information in our single cell dataset to recover regulatory interactions of lower expression TFs, although this limitation remains. Other potential areas of improvement to RENIN include including TF cooperativity and regulatory element interaction, adding any available proteomic data rather than relying on RNA as a proxy, and robust simulation capacity to further facilitate hypothesis generation.

With just two sequencing modalities, our modeling explained a large portion of variance in the expression of H-FR genes of interest. We have tailored our approach so that the output is highly interpretable in order to facilitate intuitive understanding of regulatory network composition and selection of candidates for biological validation. In agreement with previous studies, we found strong evidence that distal regulatory interactions play significant roles in regulating gene expression (6), which RENIN could leverage accurately to characterize gene regulatory networks with high resolution. This work demonstrates that increased profiling resolution can be derived from integration of different data streams, including gene expression, chromatin accessibility, and GWAS. In the future ranking CREs by predicted regulatory influence may be integrated with GWAS to increase risk variant detection and elucidate new disease mechanisms. Spatiotemporal control of transcription is achieved through layers of regulatory control that tune the accessibility of CREs through, for example, histone modifications that reduce DNA-histone interacting and facilitate DNA access by transcription factors or bind other chromatin remodeling proteins(59). Other epigenetic modifications, such as DNA methylation, may alter the ability of transcription factors to bind their cognate motifs(60). Future work on gene regulatory networks will benefit greatly from single cell profiling of epigenetic features such as methylation and histone modifications (4), in addition to the sequence information used in this study. Other improvements to modeling may include efforts to model translational regulation or non-steady-state RNA dynamics (61). The framework we have described with RENIN, in which correlated accessibility or other epigenetic marks with target gene expression, can be adapted to study the effects of these other mechanisms on gene regulation in order to construct a comprehensive model of transcriptional regulation.

## Conclusions

RENIN is a powerful tool to fully maximize the potential of investigators’ multiomic datasets. It accurately identifies key cis-regulatory elements and transcription factors in disease processes. Its utility is demonstrated here by applying it to a single nucleus multiome dataset prepared from adult human kidney samples, where it recapitulated known biology and could identify novel transcription factors underpinning a failed epithelial repair process for biological validation. Code and documentation are publicly available under the MIT license at https://www.github.com/nledru/renin.

## Methods

### Tissue Procurement

Non-tumor kidney cortex samples were obtained from patients undergoing partial or radical nephrectomy for renal mass at Brigham and Women’s Hospital (Boston, MA) under an established Institutional Review Board protocol approved by the Mass General Brigham Human Research Committee. Additional kidney cortex samples were obtained from living kidney donors prior to surgical implantation at Wake Forest University under an established protocol approved by the Wake Forest University Institutional Review Board. All participants provided written informed consent in accordance with the Declaration of Helsinki. Additional samples from kidneys rejected for transplantation were obtained from deceased organ donors provided by Mid America Transplant under an established protocol approved by the Washington University Institutional Review Board. All samples were deidentified. Samples were frozen for storage.

### Nuclear dissociation and library preparation

For single nucleus multiomic RNA- and ATAC-seq, samples were first cut into <2 mm pieces and homogenized using Dounce homogenizers and the loose head pestle (885302-0002; Kimble Chase) in 2 ml of Nuclei EZ Lysis buffer (NUC-101; Sigma-Aldrich) with protease inhibitor (5892791001; Roche) at 4°C. Samples were then filtered through a 200μm cell strainer (43-50200; pluriSelect) and homogenized in the Dounce homogenizers with the tight head pestle. Samples were incubated on ice for 5 minutes in 4 ml of EZ Lysis buffer, then filtered through a 40μm cell strainer (43-50040; pluriSelect), centrifuged at 500g for 5 minutes at 4°C. The resuspended pellet was then washed with 4ml of lysis buffer and incubated for 5 minutes at 4°C. Then the sample was centrifuged again and resuspended in Diluted Nuclei Buffer (PN-2000153; 10X Genomics), then filtered through a 5μm cell strainer (43-50005; pluriSelect). After counting, nuclei suspensions were diluted if needed to target 10,000 nuclei per lane and loaded into a thermal cycler to begin the transposition reaction, following 10X Genomics’ protocol. The manufacturer protocol was followed for the completion of library preparation.

Paired single nucleus RNA and ATAC libraries were prepared from biopsies with a modification of the protocol detailed above. Biopsies were homogenized as above, incubated for 5 minutes, filtered through a 40μm cell strainer, then spun down at 500g for 5 minutes at 4°C. The pellet was then resuspended in Diluted Nuclei Buffer and strained through a 5μm cell strainer. An aliquot of the resulting nuclei suspension was used to construct the snATAC-seq library following 10X Genomics’ protocol. The remainder was diluted with 1x DPBS (14190144; Gibco – Thermo Fisher) and used for snRNA-seq library construction following 10X Genomics’ protocol.

### Multiomic sequencing bioinformatics workflow

Seven single nucleus multiomic libraries were generated from 5 control human nephrectomy samples and 2 pre-transplant human biopsies. Libraries were sequenced on an Illumina Novaseq platform with 28-10-10-150 bp configuration for RNA-seq libraries. ATAC-seq libraries were sequenced with either a 50-8-16-50 bp configuration (1-27Nx and 2-15Nx) or 2×150 bp configuration (rest of samples). Libraries were counted with cellranger-arc 2.0.0 using the 10X-provided GRCh38-2020-A-2.0.0 reference genome and were aggregated for each sample with cellranger-arc aggr. The variational autoencoder CellBender 0.2.0 was used to reduce ambient RNA signal in each sample. The expected number of cells for each sample was estimated from cellranger-arc output, and cell probabilities were calculated for approximately 10,000 additional barcodes per sample. CellBender was run using with the following parameters: --epochs 150, --z-dim 100, --z-layers 500, -- learning-rate 0.0001. For two samples, b6 and b8, the learning rate was reduced to 0.00005 to reduce training instability. Prior to doublet removal, ArchR 1.0.1 was used for preliminary filtering (TSSEnrichment >= 4, BlacklistRatio <= 0.01, NucleosomeRatio <= 4, nFrags >= 3000, and nFrags <= 100000). Seurat 4.0.2 was also used for preliminary filtering (nCount_ATAC > 3000, nCount_ATAC < 100000, nFeature_ATAC > 1000, nCount_RNA < 50000, nCount_RNA > 1000, nFeature_RNA > 500, percent.mt < 5, percent.rps < 2, percent.rpl < 2), and cell barcodes passing both sets of filtering criteria were retained.

After the removal of low-quality barcodes, predicted doublets, and remaining small likely-doublet clusters, SCTransform was used for normalization of the snRNA-seq component, and Harmony was used for batch effect correction on the SCT output assay. To process the snATAC-seq component, term-frequency inverse-document-frequency (TFIDF) was computed, then dimensional reduction was performed on the TFIDF matrix with singular value decomposition. Harmony was used for batch effect correction on the resulting latent semantic indexing reduction. A gene activity matrix was computed with the GeneActivity function, including the 2000 bases upstream of the TSS. Cell type-specific peaks were called with MACS2 (v2.2.7.1) using the Signac CallPeaks function with default parameters for CRE analysis. A weighted shared nearest neighbor graph (SNN) was constructed on the RNA and ATAC batch-corrected dimensional reductions. Clustering of the weighted SNN graph was performed with smart local moving. A WNN UMAP reduction was calculated with the RunUMAP function for visualization.

### Doublet detection algorithm comparison

The three algorithms tested were AMULET v1.1, ArchR v1.0.1, and DoubletFinder v2.0.3. Output metadata of cellranger-arc count was modified for compatibility with AMULET with a custom script. AMULET was run on the ATAC modality for each sample, omitting genomic regions in the ENCODE hg38 blacklist(62). Barcodes with q<0.05 were identified as doublets. For doublet removal with ArchR, default settings of the addDoubletScores function are to perform 5 trials of simulating a number of doublets equal to the number of cells in the sample. ArchR’s addDoubletScores function was modified to use previously generated paired lists of randomly sampled barcodes, then performed. DoubletFinder was modified to use the same paired list of randomly sampled barcodes.

### Biopsy versus nephrectomy comparison

We aggregated our multiomic library with previously generated single nucleus RNA-seq and ATAC-seq libraries from 5 control nephrectomies (GEO accession number <u>GSE151302</u>), as well as snRNA-seq libraries generated from 3 pre-transplant biopsies and 2 pre-transplant biopsy-derived snATAC libraries. The 3 snRNA-seq libraries were processed in a similar manner to the multiome dataset, removing low quality cells with the following filtering parameters: nFeature_RNA > 500, nFeature_RNA < 6000, nCount_RNA < 16000, percent.mt < 0.8, percent.rps < 0.4, percent.rpl < 0.4. For the ATAC modalities, we aggregated fragments in the non-multiome samples into the 193,787 accessible peaks called on the multiome samples. We treated the multiome cell type annotations as ground truth cell type identities and performed label transferring to the single modality snRNA-seq and snATAC-seq datasets in order to generate unified complete datasets. We retained all nuclei with > 0.5 maximum prediction score, then used Seurat’s FindMarkers function to identify RNA and ATAC cell type markers. We then split each dataset into subsets of cell types by origin—pre-transplant biopsy or nephrectomy.

### Gene regulatory network modeling

Nuclei are first aggregated into pseudocells using a modified version of VISION’s micropooling algorithm(63). Briefly, the WNN UMAP graph is used to perform Louvain clustering, then nuclei are partitioned into pseudocells with k-means, targeting a maximum of 100 nuclei for peak aggregation, due to the increased sparsity of snATAC-seq data, and 10 nuclei for RNA aggregation. Peak accessibility and RNA expression are then averaged within pseudocells. We use these pseudocell matrices to perform two regression steps. In the first, accessible peaks within 500kbp of each modeled gene are identified as putative CREs. An adaptive elastic-net model is trained to predict the gene’s expression with peak accessibility in order to select for CREs regulating each modeled gene. Model-predicted CREs and the gene promoter region are then scanned for motifs in chromVAR’s filtered version of the cisBP v0.2 database (16,52). TFs predicted to bind a significant CRE or a peak within the gene’s promoter region (−2000 to 0 bp from TSS) are aggregated as putative regulators of each modeled gene. A second adaptive elastic-net model is then trained for each gene, using the expression of TFs with potential to bind somewhere in the CRE or promoter set. In each adaptive elastic-net step, we model the response vector *y* = (*y*_1,_ *y*_2_, … *y*_*n*_)^*T*^, where *y*_*i*_ is the expression of the target gene, *y*, in pseudocell *i* ∈ {1,2, …*n*} and *n* is the number of pseudocells, as *y* = Xβ + ϵ. The predictor matrix ***X*** contains *n* rows, for each pseudocell *i x =* (1,***x***_1_, …***x***_*p*_), where *p* is the number of regulatory elements, CREs for the first step and TFs for the second step, used to predict target gene expression. Then, the following optimization problem is solved:

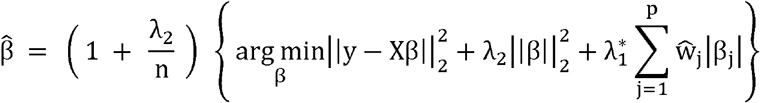

The adaptive elastic-net estimator, 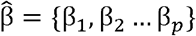, is computed for all target genes, assigning predicted weights to all included regulatory elements for each gene, with the adaptive weights, 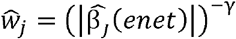 with first calculated with the elastic-net estimator(64), resulting in higher penalties for regulatory elements with lower first estimated weights. Cross-validation is used to determine the optimal *λ*_1_ values for the elastic-net and adaptive elastic-net estimators. For increasing desired levels of sparsity, *λ*_2_, can be determined by cross-validation or manually; a value of 0.5 appeared to work well as a default starting value. This estimator has the benefits of both the adaptive weighted L1 penalty and the elastic-net penalty: increasing model sparsity, producing more interpretable gene regulatory networks with fewer factors implicated, while maintaining the oracle property(22). It also allows us to simulate the effects of knocking down or upregulating select TFs, *X*_*perturb*_ as 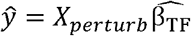. In order to determine significance, we model 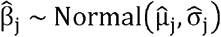. We bootstrap 1000 and 100 training sets for RENIN’s first and second steps, respectively, to estimate standard errors of each estimated regulatory weights, and retain those predicted to be nonzero with p < 0.05. Runtime with these parameters was under an hour on our dataset with multithreading.

### Cell culture

Human primary proximal tubular cells (hRPTECs; Lonza, CC-2553) were cultured with renal epithelial cell growth medium (REGM; Lonza, CC-3190). Cell cultures were maintained in humidified 5% CO_2_ at 37°C.

### CUT&RUN sequencing of RPTECs

CUT&RUN on primary RPTEC culture was performed with CUTANA kit (EpiCypher, 14-1048) according to the manufacturer’s instructions. The primary RPTEC with early passages were seeded at 8□×□10^5^ - 8□×□10^6^ cells per 10 cm culture dish 24□h prior to CUT&RUN assay. 37% formaldehyde (Sigma-Aldrich, 25259) was directly added to the medium of the RPTEC to achieve a final concentration of 0.5%, and then the medium in the dish was swirled and incubated for 1 min in room temperature. Fixation reaction was quenched by adding glycine to a final concentration of 125 mM. Subsequently, the cells were scraped from culture dishes and centrifuged at 500□×□g for 5□min. Pellets were resuspended in PBS with 1% BSA and counted. The cells were centrifuged at 500□×□g for 5□min, and resuspended with wash buffer. 500,000 cells in 100 ul wash buffer were mixed and incubated with Concanavalin A (ConA) conjugated paramagnetic beads. Antibodies were added to each sample (0.5□μg of H3K27ac antibody [Epicypher, 13-0045, 1:50], H3K4me3 antibody [Epicypher, 13-0041, 1:50], or rabbit IgG negative control antibody [Epicypher, 13-0042, 1:50]). The remaining steps were performed according to the manufacturer’s instructions for cross-linked samples. Library preparation was performed using the NEBNext Ultra II DNA Library Prep Kit for Illumina (New England BioLabs, E7645S) with the manufacturer’s instructions, including minor modifications indicated by CUTANA described above. CUT&RUN libraries were sequenced on a NovaSeq instrument (Illumina, 150□bp paired-end reads). Fastq files were trimmed with Trim Galore (Cutadapt [v2.8]) and aligned with Bowtie2 [v2.3.5.1] (parameters: --local --very-sensitive-local --no-unal --no-mixed --no-discordant --phred33 -I 10 -X 700) using hg38. Peak calling was performed using MACS2 [v2.2.7.1] with default parameters using samtools (1.9) and DeepTools (3.5.0).

### Partitioning heritability analysis of CREs

CREs that met the *p*-value threshold of 0.05 were sorted by the absolute value of their total predicted regulatory score, then binned into tertile peak sets. Bed files for each peak set were converted to hg19 with UCSC’s liftOver. The LDSC workflow was followed to partition heritability into each CRE set using the 1000G Phase 3 reference(65). GWAS summary statistics for eGFR and CKD were downloaded from the publicly available CKDGen database and formatted with munge_sumstats.py(42).

### Jaccard index

The Jaccard index is a measure of similarity between sets. It is computed as the intersection of the two sets divided by the union:

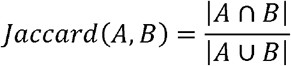

Values range from 0 in the case of no overlap to 1 in the case of perfect overlap.

### Akaike information Criterion

The Akaike Information Criterion (AIC) can be used to evaluate model goodness of fit while penalizing overfitting(66). We adapt it by calculating 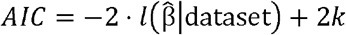 where *k* is the number of nonzero parameters for the given gene regulatory model and *l* is the log-likihood, calculated by summing 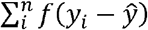 where *f* is the probability density function of the normal distribution. A smaller AIC value indicates a model with higher likelihood for a given dataset, however absolute values of AIC are meaningless without a comparison value (67). Therefore we calculate relative likelihoods between CCAN and promoter models by calculating Δ*AIC* = *AIC*_*CCAN*_ − *AIC*_*Prom*,_ with more negative values indicating a larger likelihood of the CCAN-based model relative to the promoter-only-based model. This value can be further converted into a relative probability, 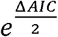.

### Motif analysis of healthy-FR PT CREs

CREs predicted to regulate differentially expressed genes between healthy and FR PT cells were identified as healthy-promoting if they positively regulated a gene upregulated in healthy PT or negatively regulated a gene upregulated in FR-PT, and failed repair-promoting if otherwise. Enriched motifs were identified with Signac’s FindMotifs for each CRE set. Footprinting analysis was performed on the subset of accessible peaks that were either healthy or FR PT CREs using a modified function adapted from Signac’s footprint functions. We downloaded a previously compiled list of hypermethylated regions associated with CKD(14), added a flanking 1 kb window, and identified overlapping peaks with findOverlaps using RENIN-predicted CREs and all 193,787 peaks called in the multiome dataset.

### Cell type-specific CRE and TF identification

The output of the two modeling steps is a set of functions of the form:

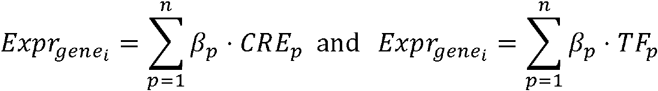

In each step, each modeled gene’s expression is the weighted sum of regulatory elements plus an intercept and error term. The weights of each regulatory element are the learned coefficients, and are a measure of that regulatory element’s influence on that gene’s expression. We take two complementary approaches to TF prioritization. The first is to rank by absolute effect. We chose to multiply each regulatory weight by the mean accessibility and expression of predicted CREs and TFs, respectively, as this would allow us to rank TFs by absolute regulatory effect on a given gene. Since this approach may be biased towards highly expressed genes, we also rank TFs by measures of centrality. After constructing a graph containing all modeled gene regulatory networks, TFs most central in the graph are prioritized as key candidate regulatory TFs. Centrality measures used for this work were PageRank and betweenness.

### siRNA knockdown of *NFAT5* in RPTECs

P2 RPTECs were switched to starvation medium for 24 hours to synchronize cell cycle. RPTECs were passaged into 6-well plates at 1*10^5^ cells and 2.5 mL of REGM per well. Fifteen uL of Lipofectamine RNAiMAX (Thermo Fisher, 13778075) was diluted in 125 uL OptiMEM (Thermo Fisher, 31985070) and 2.5 uL of ON-TARGETplus SMARTpool siRNA (Horizon Discovery, L-009618-00 for *NFAT5* targeting and D-001810-10 for non-targeting negative control pool) 50uM stock was diluted in 125 uL. The diluted siRNA and diluted Lipofectamine RNAiMAX were combined and incubated for 5 minutes, then 250 ul of the siRNA-lipid complex solution was added each well. Medium was replaced at 24 hours post-transfection, then at 48 hours, cells were harvested.

### Quantitative PCR

RNA was extracted from hRPTECs with RNeasy Mini Kit (Qiagen, 74104). Reverse transcription was performed on the RNA with High Capacity cDNA Reverse Transcription Kit (Thermo Fisher, 4368813) to prepare cDNA libraries. The Bio-Rad CFX96 Real-Time System was used for quantitative PCR, using the iTaq Universal SYBR Green Supermix (Bio-Rad). Expression of target genes were normalized to *GAPDH* expression, and the 2^−ΔΔCt^ method was used to analyze results. A two-sample t-test was performed to compare non-targeting and siRNA treatment groups, with a *p*-value < 0.05 determined to be statistically significant. Data are presented as mean ± standard deviation. Primer sequences are listed in Supplemental Table 2.

### RNA sequencing

Total RNA integrity of extracted RNA was determined by Agilent 4200 Tapestation. Library preparation was performed with 0.5 to 1 ug of total RNA. Ribosomal RNA was removed using RiboErase kits (Kapa Biosystems), then mRNA was fragmented in reverse transcriptase buffer, heating to 94° for 8 minutes.

Reverse transcription was performed with SuperScript III RT enzyme (Life Technologies) following manufacturer instructions. Illumina sequencing adapters were ligated, dual index tags were incorporated, and fragments were sequenced on an Illumina NovaSeq 6000 using paired end reads extending 150 bases. Base calls and demultiplexing were performed with Illumina’s bcl2fastq and a custom Python demultiplexing program with a maximum of one mismatch in the indexing read. Reads were aligned to the Ensembl release 101 primary assembly with STAR v2.7.9a(68) and gene counts were calculated from the number of uniquely aligned, unambiguous reads by Subread:featureCount v2.0.3(69). For all gene counts, TMM normalization size factors were calculated to adjust for differences in library size with EdgeR(70).

### NFAT5 immunofluorescence

Non-tumor human kidney cortex sample was obtained from a 71 year-old female donor (serum Cr 0.60 mg/dl). Human kidney cortex was fixed with 10% formalin overnight, embedded in paraffin, and cut at 5-μm thicknesses. Antigen retrieval in antigen unmasking solution (Vector Laboratories, H-3301-250) using was performed before staining. The sections were blocked with 1% bovine serum albumin in PBS(−) for 60 minutes at room temperature, followed by incubation with primary antibodies for NFAT5 (Thermo Fisher Scientific, PA1-023) and biotinylated Lotus Tetragonolobus Lectin (LTL) (Vector Laboratories, B-1325) at 4°C overnight. Next, the sections were incubated with secondary antibodies for 90 minutes at room temperature. Nuclei were counterstained with DAPI (4,6-diamidino-2-phenylindole) and mounted in ProLong gold antifade mountant (Thermo Fisher Scientific, P36930). Fluorescence images were captured by Nikon C2+ Eclipse confocal microscopy.

### Statistics and reproducibility

Statistical analysis was performed with publicly available code to increase reproducibility. No statistical method was used to predetermine sample size, and no data were excluded from analysis. Experiments were not randomized, and investigators were not blinded during experiments or outcome assessment.

## Declarations

This research complies with all relevant ethical regulations and has been approved by the Washington University Institutional Review Board.

## Consent for publication

Not applicable

## Availability of data and materials

The datasets generated and analyzed during the current study are available in GEO, accession number <u>GSE220289</u>. Previously generated datasets that were analyzed during the current study are available in GEO, accession number <u>GSE151302</u>. Analysis and package code are available in public repositories at https://www.github.com/nledru/renin.

## Author contributions

N.L. and B.D.H. conceived, coordinated, and designed the study. N.L., Y.M., Y.Y. and H.W. performed experiments. N.L. and P.C.W. analyzed data. S.S.W., A.A. and G.O. provided human samples. N.L. and B.D.H. wrote the manuscript. All authors read and approved the final manuscript.

## Acknowledgements

The authors acknowledge the Washington University Genome Technology Access Center and Center for Genome Sciences & Systems Biology for sequencing support.

## References

1. Lee TI, Young RA. Transcriptional Regulation and Its Misregulation in Disease. Cell. 2013 Mar;152(6):1237–51.

2. Soutourina J. Transcription regulation by the Mediator complex. Nat Rev Mol Cell Biol. 2018 Apr;19(4):262–74.

3. Oudelaar AM, Higgs DR. The relationship between genome structure and function. Nat Rev Genet. 2021 Mar;22(3):154–68.

4. Preissl S, Gaulton KJ, Ren B. Characterizing cis-regulatory elements using single-cell epigenomics. Nat Rev Genet. 2022 Jul 15; Available from: https://www.nature.com/articles/s41576-022-00509-1

5. Maurano MT, Humbert R, Rynes E, Thurman RE, Haugen E, Wang H, et al. Systematic Localization of Common Disease-Associated Variation in Regulatory DNA. Science. 2012 Sep 7;337(6099):1190–5.

6. Sieber KB, Batorsky A, Siebenthall K, Hudkins KL, Vierstra JD, Sullivan S, et al. Integrated Functional Genomic Analysis Enables Annotation of Kidney Genome-Wide Association Study Loci. J Am Soc Nephrol. 2019 Mar;30(3):421–41.

7. Wilflingseder J, Willi M, Lee HK, Olauson H, Jankowski J, Ichimura T, et al. Enhancer and super-enhancer dynamics in repair after ischemic acute kidney injury. Nat Commun. 2020 Dec;11(1):3383.

8. Wilson PC, Wu H, Kirita Y, Uchimura K, Ledru N, Rennke HG, et al. The single-cell transcriptomic landscape of early human diabetic nephropathy. Proc Natl Acad Sci. 2019 Sep 24;116(39):19619–25.

9. Kirita Y, Wu H, Uchimura K, Wilson PC, Humphreys BD. Cell profiling of mouse acute kidney injury reveals conserved cellular responses to injury. Proc Natl Acad Sci U S A. 2020 07;117(27):15874–83.

10. Gerhardt LMS, Liu J, Koppitch K, Cippà PE, McMahon AP. Single-nuclear transcriptomics reveals diversity of proximal tubule cell states in a dynamic response to acute kidney injury. Proc Natl Acad Sci. 2021 Jul 6;118(27):e2026684118.

11. Kuppe C, Ibrahim MM, Kranz J, Zhang X, Ziegler S, Perales-Patón J, et al. Decoding myofibroblast origins in human kidney fibrosis. Nature. 2020 Nov 11; Available from: http://www.nature.com/articles/s41586-020-2941-1

12. Muto Y, Dixon EE, Yoshimura Y, Wu H, Omachi K, Ledru N, et al. Defining cellular complexity in human autosomal dominant polycystic kidney disease by multimodal single cell analysis. Nat Commun. 2022 Oct 30;13(1):6497.

13. Wu H, Gonzalez Villalobos R, Yao X, Reilly D, Chen T, Rankin M, et al. Mapping the single-cell transcriptomic response of murine diabetic kidney disease to therapies. Cell Metab. 2022 Jul;34(7):1064-1078.e6.

14. Wilson PC, Muto Y, Wu H, Karihaloo A, Waikar SS, Humphreys BD. Multimodal single cell sequencing implicates chromatin accessibility and genetic background in diabetic kidney disease progression. Nat Commun. 2022 Sep 6;13(1):5253.

15. Xu Y, Kuppe C, Perales-Patón J, Hayat S, Kranz J, Abdallah AT, et al. Adult human kidney organoids originate from CD24+ cells and represent an advanced model for adult polycystic kidney disease. Nat Genet. 2022 Nov;54(11):1690–701.

16. Weirauch MT, Yang A, Albu M, Cote AG, Montenegro-Montero A, Drewe P, et al. Determination and Inference of Eukaryotic Transcription Factor Sequence Specificity. Cell. 2014 Sep;158(6):1431–43.

17. Aibar S, González-Blas CB, Moerman T, Huynh-Thu VA, Imrichova H, Hulselmans G, et al. SCENIC: single-cell regulatory network inference and clustering. Nat Methods. 2017 Nov;14(11):1083–6.

18. Smemo S, Tena JJ, Kim KH, Gamazon ER, Sakabe NJ, Gómez-Marín C, et al. Obesity-associated variants within FTO form long-range functional connections with IRX3. Nature. 2014 Mar 20;507(7492):371–5.

19. Thurman RE, Rynes E, Humbert R, Vierstra J, Maurano MT, Haugen E, et al. The accessible chromatin landscape of the human genome. Nature. 2012 Sep;489(7414):75–82.

20. Ma S, Zhang B, LaFave L, Chiang Z, Hu Y, Ding J, et al. Chromatin potential identified by shared single cell profiling of RNA and chromatin. Genomics; 2020 Jun. Available from: http://biorxiv.org/lookup/doi/10.1101/2020.06.17.156943

21. Zhang L, Zhang J, Nie Q. DIRECT-NET: An efficient method to discover cis-regulatory elements and construct regulatory networks from single-cell multiomics data. Sci Adv. 2022 Jun 3;8(22):eabl7393.

22. Zou H, Zhang HH. On the adaptive elastic-net with a diverging number of parameters. Ann Stat. 2009 Aug 1;37(4). Available from: https://projecteuclid.org/journals/annals-of-statistics/volume-37/issue-4/On-the-adaptive-elastic-net-with-a-diverging-number-of/10.1214/08-AOS625.full

23. Chang-Panesso M, Kadyrov FF, Lalli M, Wu H, Ikeda S, Kefaloyianni E, et al. FOXM1 drives proximal tubule proliferation during repair from acute ischemic kidney injury. J Clin Invest. 2019 Nov 11;129(12):5501–17.

24. Ichimura T, Asseldonk EJP v., Humphreys BD, Gunaratnam L, Duffield JS, Bonventre JV. Kidney injury molecule–1 is a phosphatidylserine receptor that confers a phagocytic phenotype on epithelial cells. J Clin Invest. 2008 May 1;118(5):1657–68.

25. Yang L, Besschetnova TY, Brooks CR, Shah JV, Bonventre JV. Epithelial cell cycle arrest in G2/M mediates kidney fibrosis after injury. Nat Med. 2010 May;16(5):535–43.

26. Hinze C, Kocks C, Leiz J, Karaiskos N, Boltengagen A, Cao S, et al. Single-cell transcriptomics reveals common epithelial response patterns in human acute kidney injury. Genome Med. 2022 Sep 9;14(1):103.

27. Muto Y, Wilson PC, Wu H, Waikar SS, Humphreys B. Single cell transcriptional and chromatin accessibility profiling redefine cellular heterogeneity in the adult human kidney. Genomics; 2020 Jun. Available from: http://biorxiv.org/lookup/doi/10.1101/2020.06.14.151167

28. Muto Y, Wilson PC, Ledru N, Wu H, Dimke H, Waikar SS, et al. Single cell transcriptional and chromatin accessibility profiling redefine cellular heterogeneity in the adult human kidney. Nat Commun. 2021 Dec;12(1):2190.

29. Fleming SJ, Marioni JC, Babadi M. CellBender remove-background: a deep generative model for unsupervised removal of background noise from scRNA-seq datasets. Bioinformatics; 2019 Oct. Available from: http://biorxiv.org/lookup/doi/10.1101/791699

30. McGinnis CS, Murrow LM, Gartner ZJ. DoubletFinder: Doublet Detection in Single-Cell RNA Sequencing Data Using Artificial Nearest Neighbors. Cell Syst. 2019 Apr;8(4):329-337.e4.

31. Thibodeau A, Eroglu A, McGinnis CS, Lawlor N, Nehar-Belaid D, Kursawe R, et al. AMULET: a novel read count-based method for effective multiplet detection from single nucleus ATAC-seq data. Genome Biol. 2021 Dec;22(1):252.

32. Korsunsky I, Millard N, Fan J, Slowikowski K, Zhang F, Wei K, et al. Fast, sensitive and accurate integration of single-cell data with Harmony. Nat Methods. 2019 Dec;16(12):1289–96.

33. Stuart T, Butler A, Hoffman P, Hafemeister C, Papalexi E, Mauck WM, et al. Comprehensive Integration of Single-Cell Data. Cell. 2019 Jun;177(7):1888-1902.e21.

34. Hao Y, Hao S, Andersen-Nissen E, Mauck WM, Zheng S, Butler A, et al. Integrated analysis of multimodal single-cell data. Cell. 2021 Jun;184(13):3573-3587.e29.

35. Granja JM, Corces MR, Pierce SE, Bagdatli ST, Choudhry H, Chang HY, et al. ArchR is a scalable software package for integrative single-cell chromatin accessibility analysis. Nat Genet. 2021 Mar;53(3):403–11.

36. Xi NM, Li JJ. Benchmarking Computational Doublet-Detection Methods for Single-Cell RNA Sequencing Data. Cell Syst. 2021 Feb;12(2):176-194.e6.

37. McEvoy CM, Murphy JM, Zhang L, Clotet-Freixas S, Mathews JA, An J, et al. Single-cell profiling of healthy human kidney reveals features of sex-based transcriptional programs and tissue-specific immunity. Nat Commun. 2022 Dec 10;13(1):7634.

38. Pliner HA, Packer JS, McFaline-Figueroa JL, Cusanovich DA, Daza RM, Aghamirzaie D, et al. Cicero Predicts cis-Regulatory DNA Interactions from Single-Cell Chromatin Accessibility Data. Mol Cell. 2018 Sep;71(5):858-871.e8.

39. Zou H. The Adaptive Lasso and Its Oracle Properties. J Am Stat Assoc. 2006 Dec 1;101(476):1418–29.

40. Skene PJ, Henikoff S. An efficient targeted nuclease strategy for high-resolution mapping of DNA binding sites. eLife. 2017 Jan 16;6:e21856.

41. Kimura H. Histone modifications for human epigenome analysis. J Hum Genet. 2013 Jul;58(7):439–45.

42. Wuttke M, Li Y, Li M, Sieber KB, Feitosa MF, Gorski M, et al. A catalog of genetic loci associated with kidney function from analyses of a million individuals. Nat Genet. 2019;51(6):957–72.

43. Guo JK. WT1 is a key regulator of podocyte function: reduced expression levels cause crescentic glomerulonephritis and mesangial sclerosis. Hum Mol Genet. 2002 Mar 1;11(6):651–9.

44. Maezawa Y, Onay T, Scott RP, Keir LS, Dimke H, Li C, et al. Loss of the Podocyte-Expressed Transcription Factor Tcf21/Pod1 Results in Podocyte Differentiation Defects and FSGS. J Am Soc Nephrol. 2014 Nov;25(11):2459–70.

45. Usui T, Morito N, Shawki HH, Sato Y, Tsukaguchi H, Hamada M, et al. Transcription factor MafB in podocytes protects against the development of focal segmental glomerulosclerosis. Kidney Int. 2020 Aug;98(2):391–403.

46. Dhillon P, Park J, Hurtado del Pozo C, Li L, Doke T, Huang S, et al. The Nuclear Receptor ESRRA Protects from Kidney Disease by Coupling Metabolism and Differentiation. Cell Metab. 2020 Dec;S1550413120306069.

47. Miao Z, Balzer MS, Ma Z, Liu H, Wu J, Shrestha R, et al. Single cell regulatory landscape of the mouse kidney highlights cellular differentiation programs and disease targets. Nat Commun. 2021 Dec;12(1):2277.

48. Ronconi E, Sagrinati C, Angelotti ML, Lazzeri E, Mazzinghi B, Ballerini L, et al. Regeneration of Glomerular Podocytes by Human Renal Progenitors. J Am Soc Nephrol. 2009 Feb;20(2):322–32.

49. Appel D, Kershaw DB, Smeets B, Yuan G, Fuss A, Frye B, et al. Recruitment of Podocytes from Glomerular Parietal Epithelial Cells. J Am Soc Nephrol. 2009 Feb;20(2):333–43.

50. Nekulova M, Holcakova J, Coates P, Vojtesek B. The role of P63 in cancer, stem cells and cancer stem cells. Cell Mol Biol Lett. 2011 Jan 1;16(2). Available from: https://www.degruyter.com/document/doi/10.2478/s11658-011-0009-9/html

51. Shankland SJ, Smeets B, Pippin JW, Moeller MJ. The emergence of the glomerular parietal epithelial cell. Nat Rev Nephrol. 2014 Mar;10(3):158–73.

52. Schep AN, Wu B, Buenrostro JD, Greenleaf WJ. chromVAR: inferring transcription-factor-associated accessibility from single-cell epigenomic data. Nat Methods. 2017 Oct;14(10):975–8.

53. Zhao J, Lupino K, Wilkins BJ, Qiu C, Liu J, Omura Y, et al. Genomic integration of ERRγ-HNF1β regulates renal bioenergetics and prevents chronic kidney disease. Proc Natl Acad Sci. 2018 May 22;115(21). Available from: https://pnas.org/doi/full/10.1073/pnas.1804965115

54. Bonomo JA, Guan M, Ng MCY, Palmer ND, Hicks PJ, Keaton JM, et al. The ras responsive transcription factor RREB1 is a novel candidate gene for type 2 diabetes associated end-stage kidney disease. Hum Mol Genet. 2014 Dec 15;23(24):6441–7.

55. Chen L, Luo S, Dupre A, Vasoya RP, Parthasarathy A, Aita R, et al. The nuclear receptor HNF4 drives a brush border gene program conserved across murine intestine, kidney, and embryonic yolk sac. Nat Commun. 2021 May 17;12(1):2886.

56. Melchinger I, Guo K, Guo J, Xu L. Inflammation-mediated Upregulation of VCAM-1 but not KIM-1 during Acute Kidney Injury to Chronic Kidney Disease Transition. Pathology; 2022 Sep. Available from: http://biorxiv.org/lookup/doi/10.1101/2022.09.15.508151

57. Hao S, Bellner L, Zhao H, Ratliff BB, Darzynkiewicz Z, Vio CP, et al. NFAT5 Is Protective Against Ischemic Acute Kidney Injury. Hypertension. 2014 Mar;63(3). Available from: https://www.ahajournals.org/doi/10.1161/HYPERTENSIONAHA.113.02476

58. Dietterich TG. An Experimental Comparison of Three Methods for Constructing Ensembles of Decision Trees: Bagging, Boosting, and Randomization. Mach Learn. 2000;40(2):139–57.

59. Bannister AJ, Kouzarides T. Regulation of chromatin by histone modifications. Cell Res. 2011 Mar;21(3):381–95.

60. Portela A, Esteller M. Epigenetic modifications and human disease. Nat Biotechnol. 2010 Oct;28(10):1057–68.

61. Stark R, Grzelak M, Hadfield J. RNA sequencing: the teenage years. Nat Rev Genet. 2019 Nov;20(11):631–56.

62. Amemiya HM, Kundaje A, Boyle AP. The ENCODE Blacklist: Identification of Problematic Regions of the Genome. Sci Rep. 2019 Dec;9(1):9354.

63. DeTomaso D, Jones MG, Subramaniam M, Ashuach T, Ye CJ, Yosef N. Functional interpretation of single cell similarity maps. Nat Commun. 2019 Dec;10(1):4376.

64. Zou H, Hastie T. Regularization and variable selection via the elastic net. J R Stat Soc Ser B Stat Methodol. 2005 Apr;67(2):301–20.

65. ReproGen Consortium, Schizophrenia Working Group of the Psychiatric Genomics Consortium, The RACI Consortium, Finucane HK, Bulik-Sullivan B, Gusev A, et al. Partitioning heritability by functional annotation using genome-wide association summary statistics. Nat Genet. 2015 Nov;47(11):1228–35.

66. Akaike H. Information Theory and an Extension of the Maximum Likelihood Principle. In: Parzen E, Tanabe K, Kitagawa G, editors. Selected Papers of Hirotugu Akaike. New York, NY: Springer New York; 1998. p. 199–213. (Springer Series in Statistics). Available from: http://link.springer.com/10.1007/978-1-4612-1694-0_15

67. Burnham KP, Anderson DR. Multimodel Inference: Understanding AIC and BIC in Model Selection. Sociol Methods Res. 2004 Nov;33(2):261–304.

68. Dobin A, Davis CA, Schlesinger F, Drenkow J, Zaleski C, Jha S, et al. STAR: ultrafast universal RNA-seq aligner. Bioinformatics. 2013 Jan;29(1):15–21.

69. Liao Y, Smyth GK, Shi W. featureCounts: an efficient general purpose program for assigning sequence reads to genomic features. Bioinformatics. 2014 Apr 1;30(7):923–30.

70. Robinson MD, McCarthy DJ, Smyth GK. edgeR: a Bioconductor package for differential expression analysis of digital gene expression data. Bioinformatics. 2010 Jan 1;26(1):139–40.

