## Supplementary Information for "Predicting regulators of epithelial cell state through regularized regression analysis of single cell multiomic sequencing"

Table of Contents

**Donor Demographics and Clinical Data**

Supplemental Table 1	3

**Primer sequences used for qPCR experiments**

Supplemental Table 2	4

**Simultaneous single-nucleus transcriptional and chromatin accessibility profiling of the adult human kidney resolves high-quality cell type-specific profiles**

Supplemental Figure 1	5

**Doublet calling algorithms identify distinct sets of doublets**

Supplemental Figure 2	6

**Partial nephrectomy kidney samples are similar to live donor samples and both have FR-PT cells**

Supplemental Figure 3	7

**Regulatory network inference with adaptive elastic-net model**

Supplemental Figure 4	8

**Cell type-specific *cis-*regulatory elements identified with RENIN**

Supplemental Figure 5	9

Supplemental Figure 6	10

**CRE analysis of healthy-failed repair axis reveals coordinated regulatory element remodeling**

Supplemental Figure 7	11

**Identification of healthy- and FR-promoting TFs with RENIN**

Supplemental Figure 8	12

***NFAT5* knockdown partially reverts failed repair phenotype**

Supplemental Figure 9	13

| **ID** | **Age** | **Race (AA = African American, W = White)** | **Sex** | **sCr (mg/dL)** | **Glomerulosclerosis** | **IFTA** | **ANS** | **Sample type** | **Modality** |
| --- | --- | --- | --- | --- | --- | --- | --- | --- | --- |
| N1 | 65 | W | M | 1.01 | <10% | Mild | Moderate-Severe | Nephrectomy | Multiome |
| N2 | 64 | W | F | 0.99 | <10% | None | None | Nephrectomy | Multiome |
| A1 | 45 | AA | F | 0.49 | <10% | Mild (1-10%) | Moderate | Nephrectomy | Multiome |
| A2 | 73 | AA | M | 4.73 | 10-20% | Moderate (10-20%) | Severe | Nephrectomy | Multiome |
| A3 | 76 | AA | F | 0.95 | 10-20% | Moderate | Moderate | Nephrectomy | Multiome |
| B6* | 46 | W | F | 2.3 | <10% | Mild | Mild | Biopsy – Living donor | Multiome |
| B7 | 58 | W | F | 0.6 | <10% | Mild | Moderate | Biopsy - DCD | Multiome |
| B3* | 71 | W | F | 0.7 | Normal biopsy | Normal biopsy | Normal biopsy | Biopsy – Living donor | Split single nucleus RNA-seq and ATAC-seq |
| B4 | 50 | W | F | 0.7 | <10% | Mild | None | Biopsy - DCD | Split single nucleus RNA-seq and ATAC-seq |
| B5 | 48 | AA | M | 1.2 | <10% | Mild | Mild | Biopsy – Living donor | Split single nucleus RNA-seq and ATAC-seq |

**Supplemental Table 1. Donor Demographics and Clinical Data.** Asterisks indicate IDs from which libraries were generated for two samples, each from contralateral kidneys. Biopsies were either from living donors or DCD, indicating donation after cardiac death.

| **Primer name** | **Sequence (5' to 3')** |
| --- | --- |
| NFAT5 F | CCTAATGCCCTGATGACTCCAC |
| NFAT5 R | GTTTGCTGAGTTGATCCAACAGAC |
| VCAM1 F | GATTCTGTGCCCACAGTAAGGC |
| VCAM1 R | TGGTCACAGAGCCACCTTCTTG |
| GAPDH F | GACAGTCAGCCGCATCTTCT |
| GAPDH R | GCGCCCAATACGACCAAATC |

**Supplemental Table 2. Primer sequences used for qPCR experiments.**

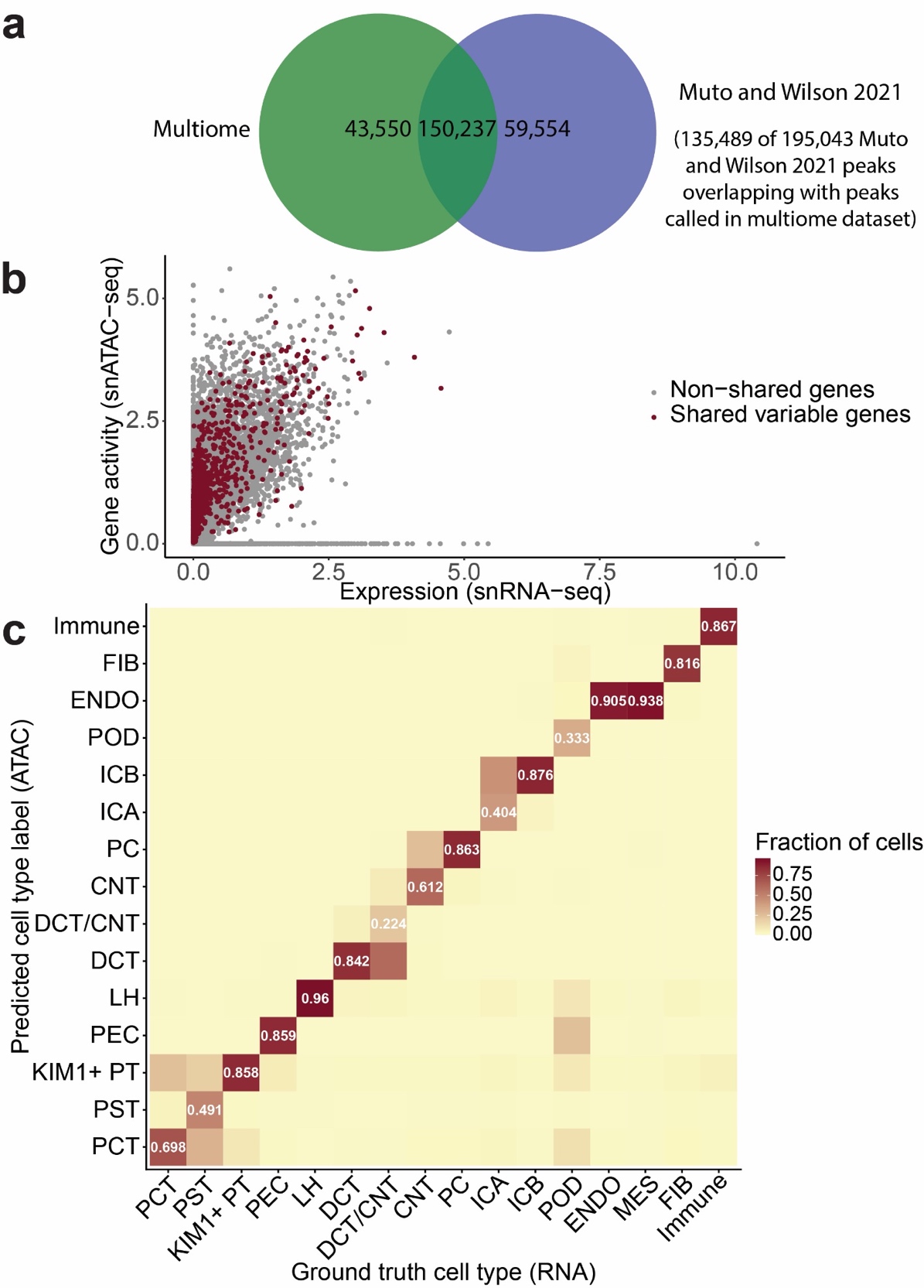

**Supplemental Figure 1. Analysis of ATAC modality in single nucleus multiomic dataset. a.** Overlap between peaks called on this dataset and prior snATAC-seq dataset generated in our lab. **b.** Gene activity scores versus expression for all genes in dataset. Genes that were selected as top variable features in both the Gene Activity and RNA assays are colored red. **c.** Confusion matrix showing agreement between cell type annotations as determined by RNA marker expression and cell type predictions using Seurat’s LabelTransfer method.

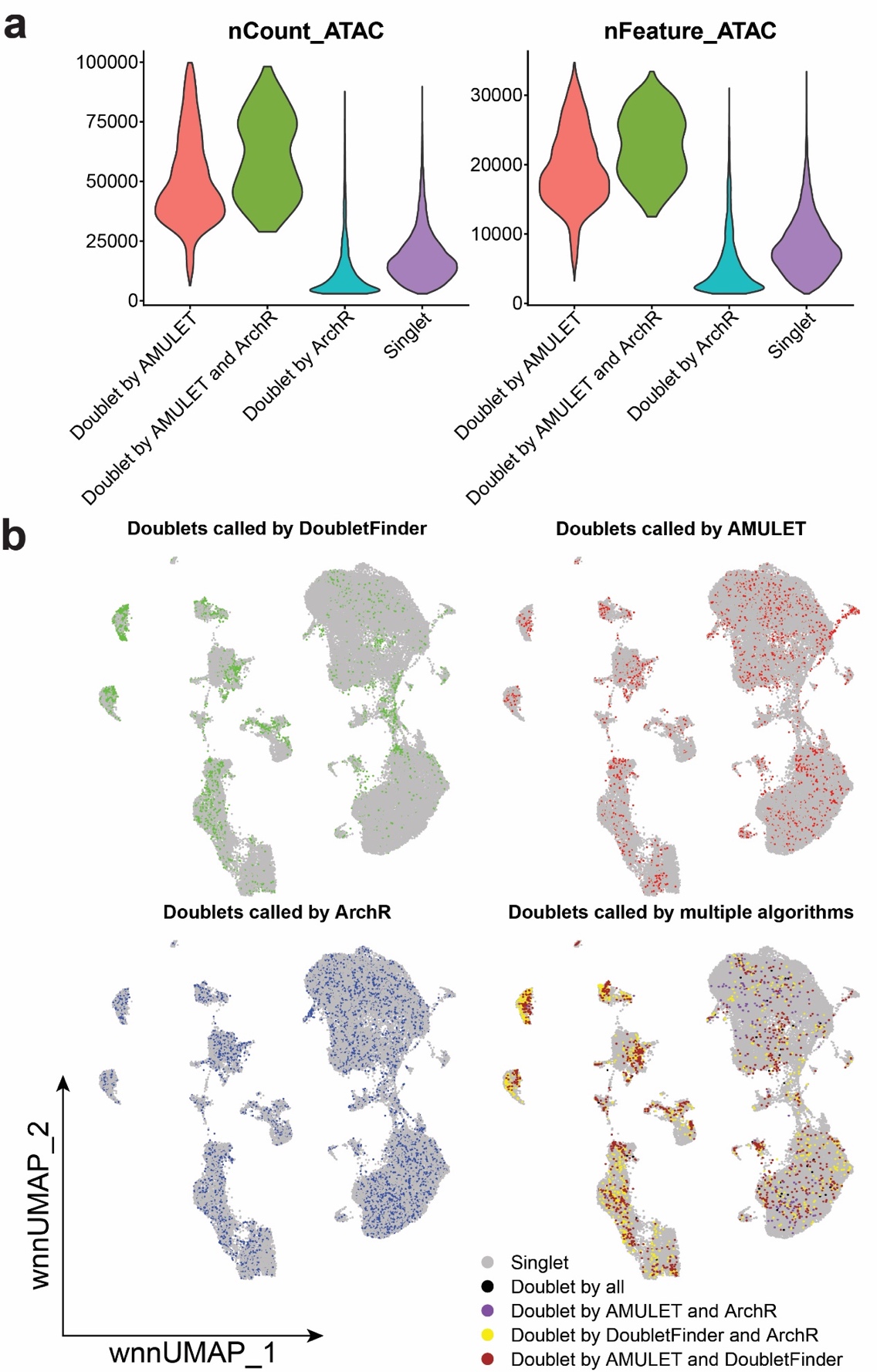

**Supplemental Figure 2. Comparison of doublet prediction algorithms. a.** Violin plots for number of unique and total peaks per barcode, labeled by doublet prediction by ArchR and AMULET. **b.** counterclockwise from top left, WNN UMAP plots of dataset with doublet predictions by AMULET, DoubletFinder, ArchR, and by all three algorithms.

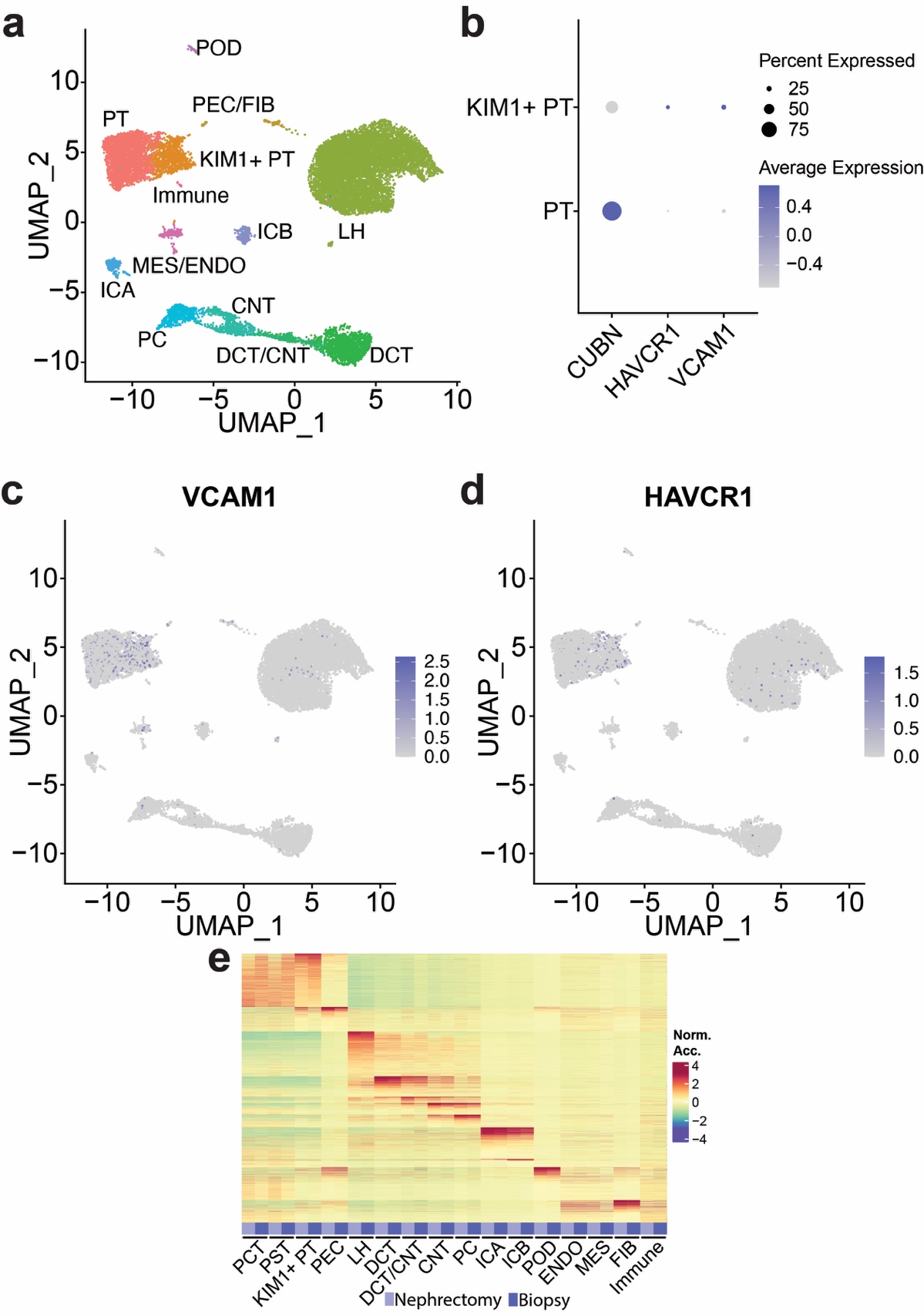

**Supplemental Figure 3. KIM1+/VCAM1+ PT population present in living donor biopsy-derived samples. a.** UMAP plot of reclustered living donor biopsy-derived-only dataset (n = 5 from 3 individual donors). **b.** Expression dot plot of healthy PT (*CUBN*) and failed repair PT (*VCAM1* and *HAVCR1*) marker genes. **c.** *VCAM1* expression and **d.** *HAVCR1* expression in living donor biopsy-only dataset. **e.** Heatmap of cell type marker peak accessibility for each cell type by sample type—nephrectomy or living donor biopsy.

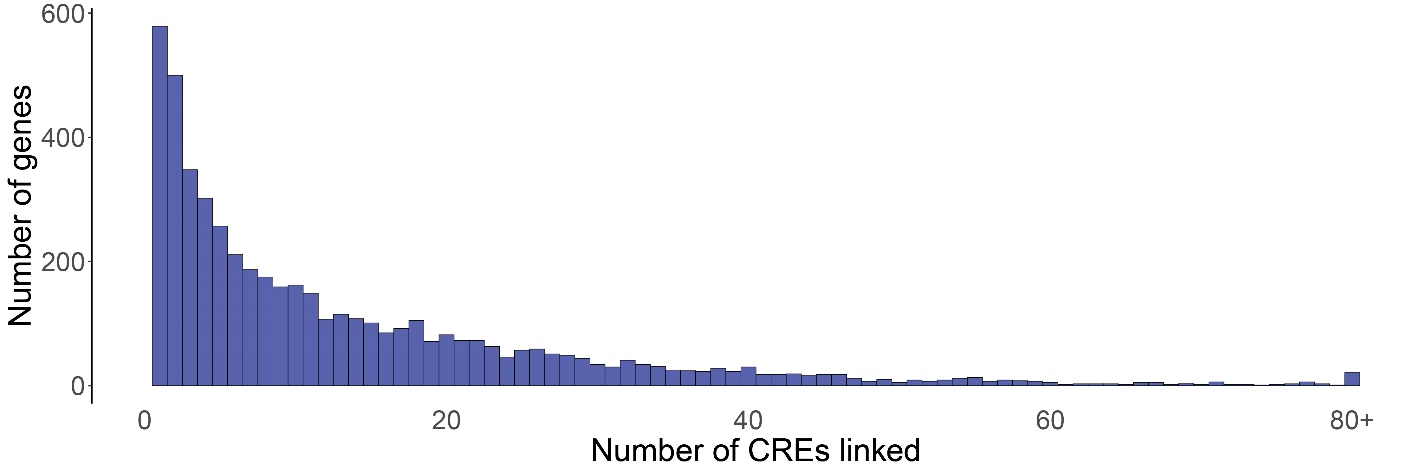

**Supplemental Figure 4. RENIN predicts multiple CREs linked to most modeled genes.** Histogram showing modeled cell type marker genes binned by number of RENIN-predicted linked CREs.

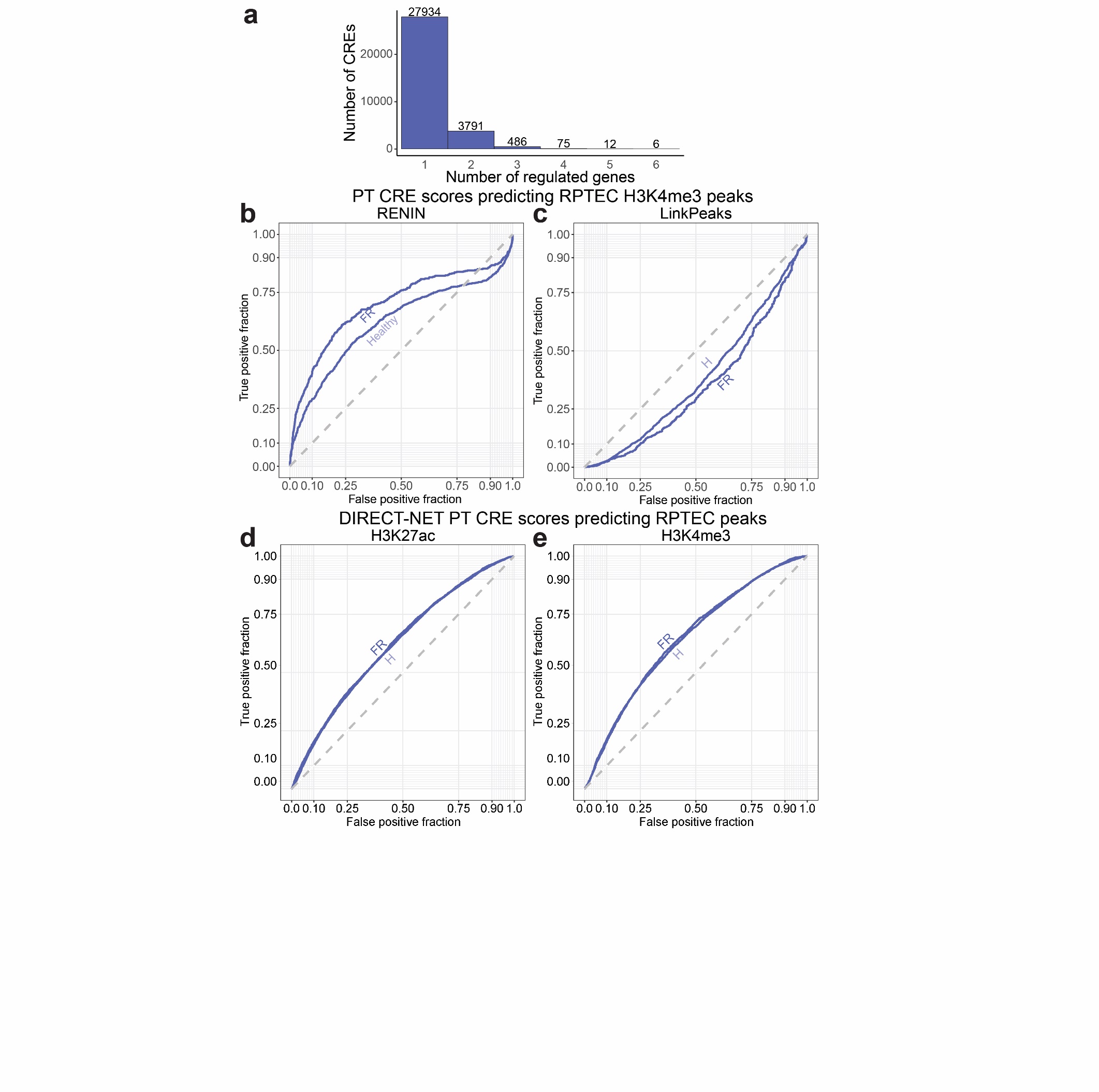

**Supplemental Figure 5. Assessment of RENIN CRE predictions relative to LinkPeaks and DIRECT-NET. a.** Histogram showing RENIN-predicted CREs binned by number of predicted linked genes. **b.** ROC curve calculated for RENIN-predicted healthy-FR PT CREs against RPTEC H3K4me3 peaks identified with CUT&RUN. AUCs were 0.696 (FR) and 0.617 (Healthy). **c.** ROC curve calculated for LinkPeaks-predicted healthy-FR PT CREs against same RPTEC H3K4me3 peak set as in (**c**). AUCs were 0.350 (FR) and 0.388 (Healthy). **d** and **e.** ROC curve calculated for DIRECT-NET-predicted healthy-FR PT CREs against RPTEC H3K27ac (AUCs 0.630 and 0.621) (**d**) and H3K4me3 (AUCs 0.649 and 0.644) (**e**) CUT&RUN peaks.

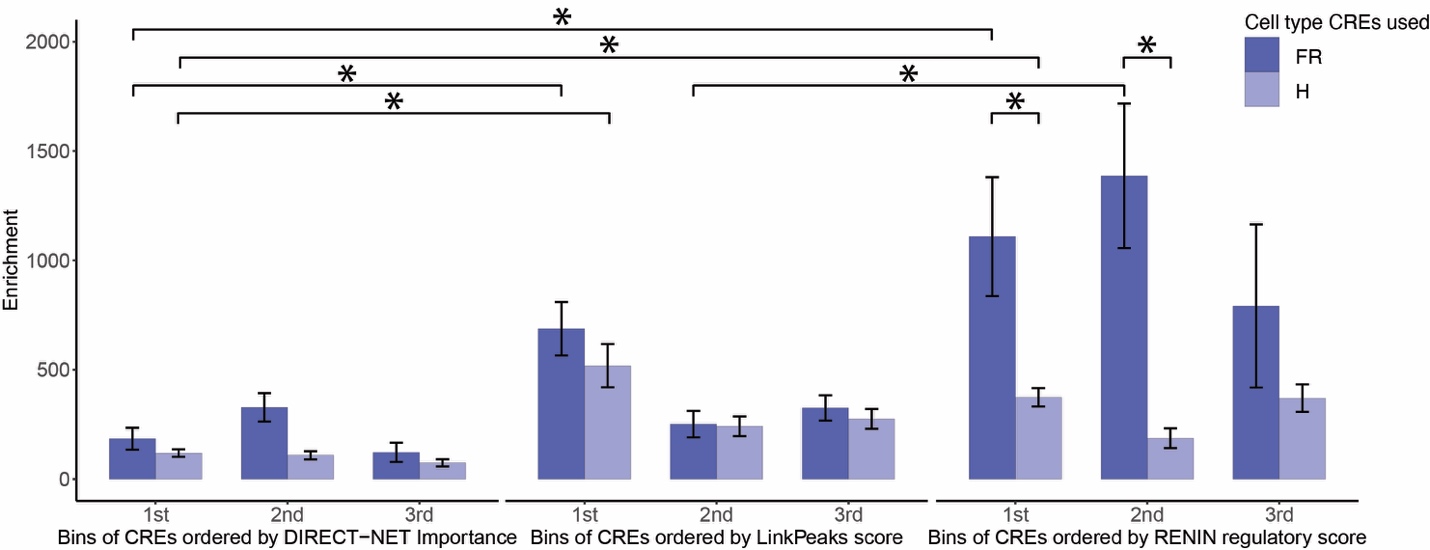

**Supplemental Figure 6. RENIN-predicted CREs are enriched for eGFR heritability.** Comparison of RENIN, LinkPeaks, and DIRECT-NET by enrichment of partitioned heritability of eGFR in model-predicted healthy (PCT + PST) and FR (failed repair—KIM1+ PT) CREs. Statistical significance determined by two-tailed t-test of difference between enrichment, p < 0.05.

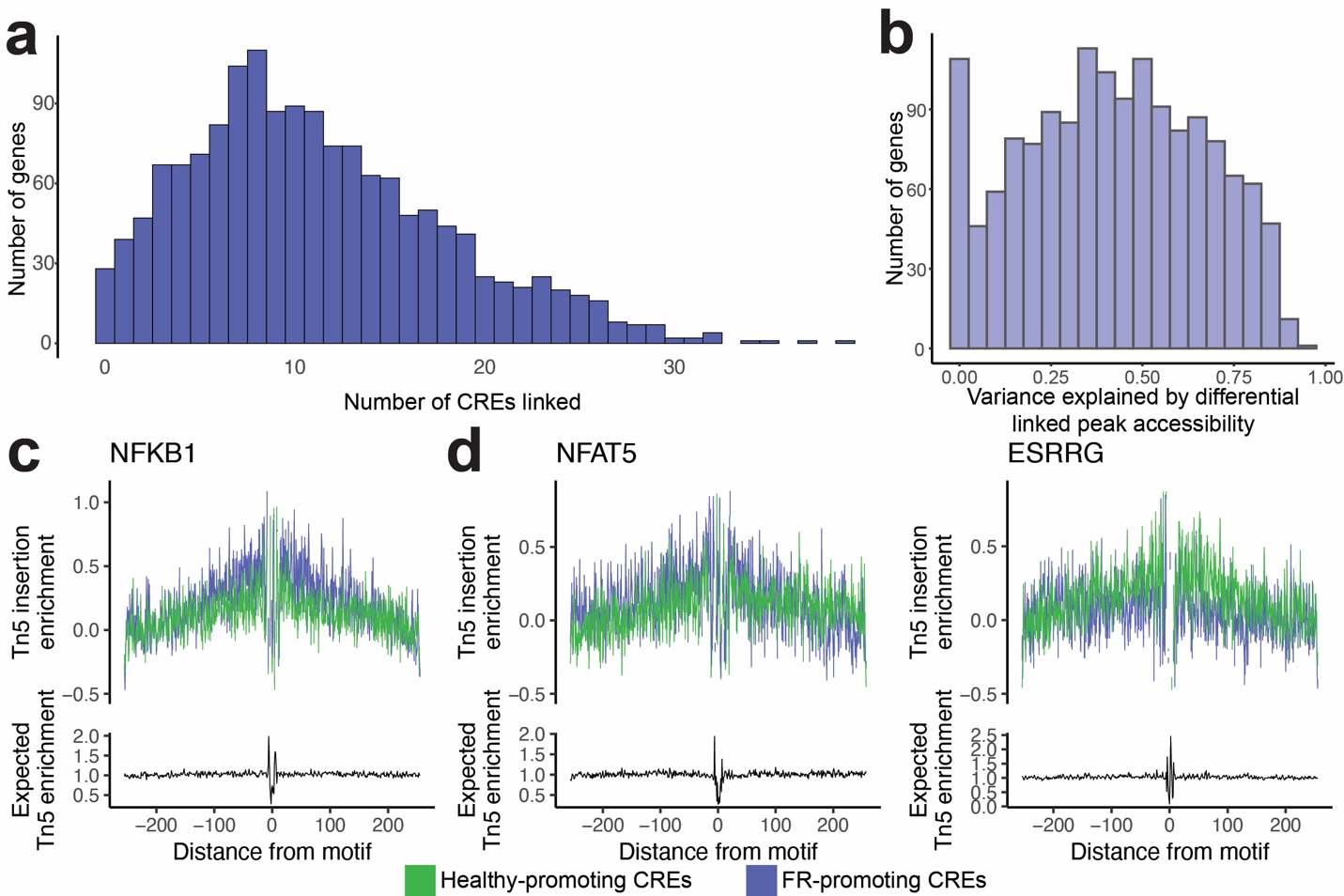

**Supplemental Figure 7. RENIN models CRE regulation of healthy-FR PT differentially expressed genes. a.** Histogram showing 1,488 healthy-FR PT modeled genes binned by the number of RENIN-predicted linked CREs. **b.** Histogram of coefficients of determination of model predictions of DEG expression versus actual expression when training model to use linked CRE accessibility as predictor variables. **c**. Footprinting analysis for NFKB1. **d.** Footprinting analysis for NFAT5, left, and ESRRG, right. Tn5 insertion enrichment calculated around motifs present in healthy-promoting CREs (green) and motifs present in FR-promoting CREs (purple).

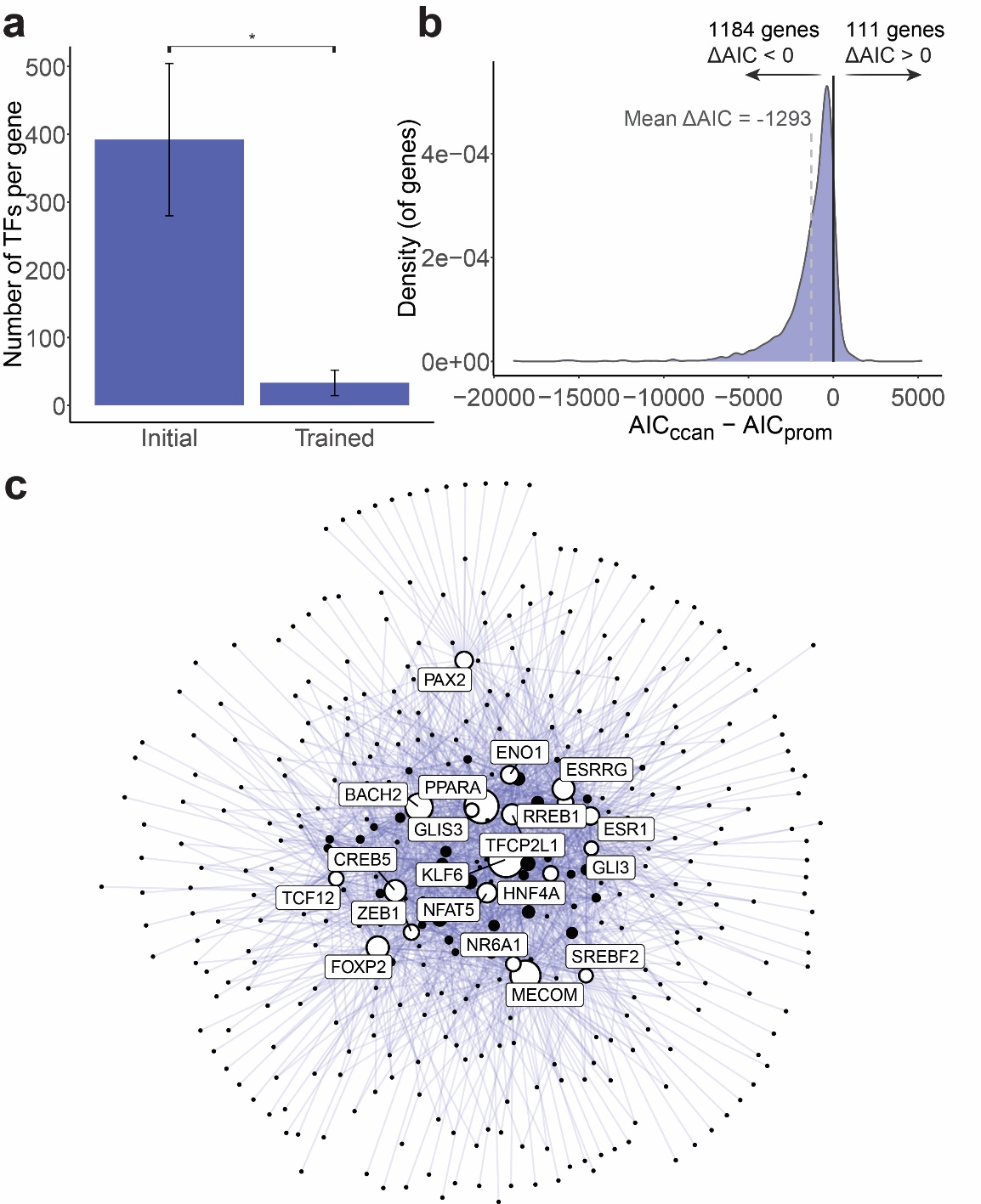

**Supplemental Figure 8. RENIN training constructs pruned and accurate gene regulatory networks. a.** Number of TFs per modeled gene before and after RENIN training. The initial number consists of all unique TF motifs present in a gene’s linked CREs and promoter peaks. The trained number of TFs consists of the number of TFs with RENIN-predicted non-zero regulatory weights for that gene. Statistical significance evaluated by paired 2-tailed t-test, p < 2.2e-16. **b.** Density plot of ΔAIC for 1323 modeled genes with at least one promoter peak (MACS2-called peak within 2000bp of TSS), calculating ΔAIC as the difference between the AIC of the RENIN-trained model using TFs within all predicted linked CREs (CCAN—*cis*-coaccessibility network) and the AIC of the trained model using TFs within the promoter peak(s) only. Mean (gray dashed line) and median ΔAIC are -1293 and -771, respectively. 1184 genes have a negative ΔAIC, while 111 have a positive ΔAIC (black line). **c**. Graph visualization of gene regulatory networks predicted by RENIN. TF node size represents centrality computed by betweenness, top 20 TFs are labeled.

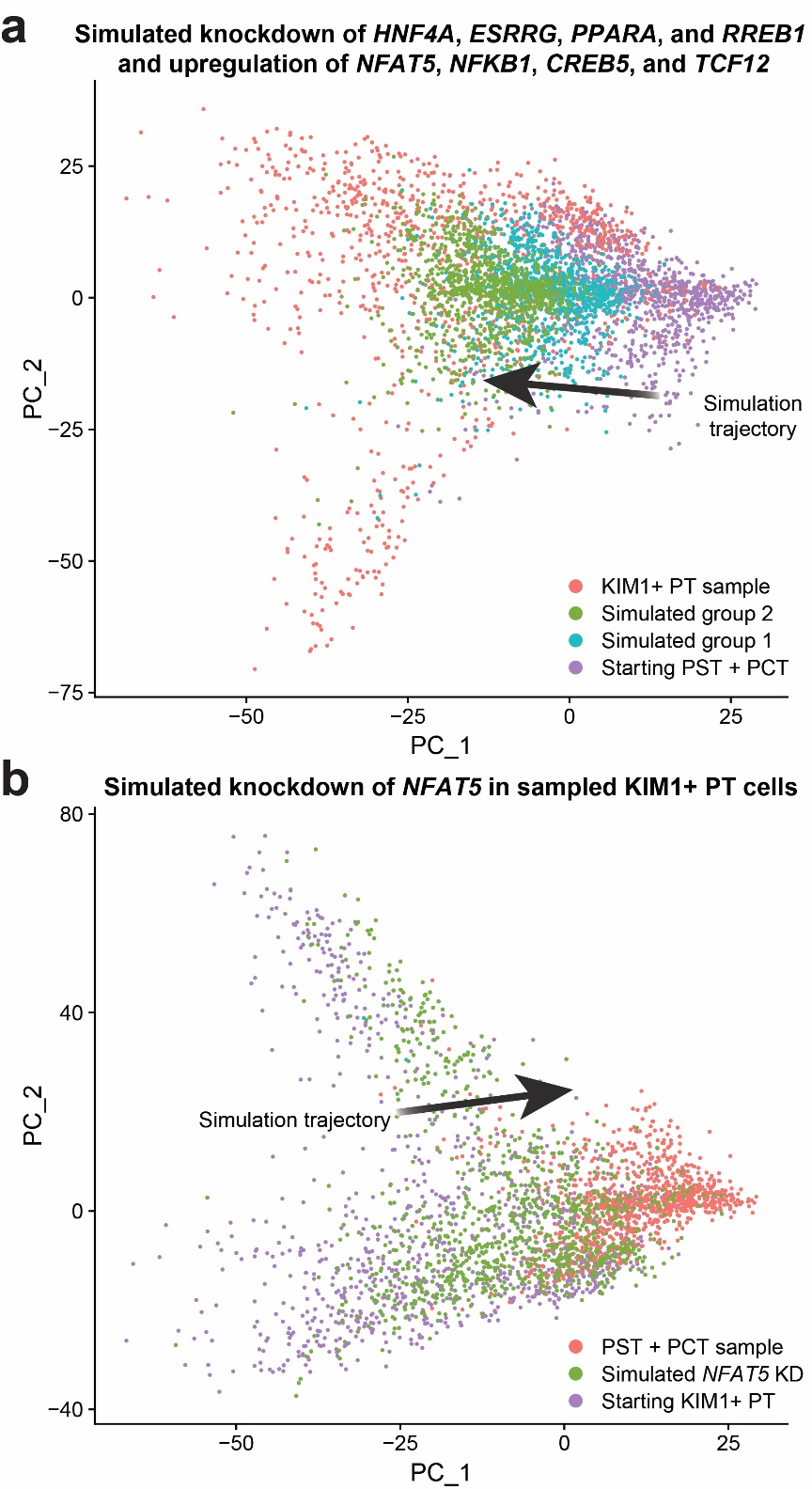

**Supplemental Figure 9. RENIN predictions can be used to simulate effects of perturbing key TFs. a.** PCA plot of subsampled PT dataset, with 1000 KIM1+ PT cells and 1000 starting PST + PCT cells. Simulated group 1 and 2 are results of upregulating *HNF4A*, *ESRRG*, *PPARA*, and *RREB1* and downregulating *NFAT5*, *NFKB1*, *CREB5*, and *TCF12* by 5 (group 1) and 10 (group 2) standard deviations in the starting PST + PCT sample, calculated by each TF’s expression in the PT dataset. Downregulation that would cause a TF’s simulated expression to be negative is set to 0. **b.** PCA plot of subsampled PT dataset, with 1000 PST and PCT cells and 1000 starting KIM1+ PT cells. *NFAT5* knockdown to an expression of 0 is simulated in these starting KIM1+ PT cells (Simulated *NFAT5* KD group).
